# Characterization and genome analysis of a psychrophilic methanotroph representing a ubiquitous *Methylobacter* spp. cluster in boreal lake ecosystems

**DOI:** 10.1101/2022.05.24.493254

**Authors:** Ramita Khanongnuch, Rahul Mangayil, Mette Marianne Svenning, Antti Juhani Rissanen

## Abstract

Lakes and ponds are considered as a major natural source of CH_4_ emissions, particularly during the ice-free period in boreal ecosystems. Aerobic methane oxidizing bacteria (MOB), which utilize CH_4_ using oxygen as an electron acceptor, are one of dominant microorganisms in the CH_4_-rich water columns. The metagenome-assembled genomes (MAGs) have revealed the genetic potential of MOB from boreal aquatic ecosystems for various microaerobic/anaerobic metabolic functions; however, the experimental validation of the process has not been succeeded. Additionally, psychrophilic (i.e., cold loving) MOB isolates and their CH_4_ oxidizing process have rarely been investigated. In this study, we isolated, provided taxonomic description, and analyzed the genome of *Methylobacte*r sp. S3L5C, a psychrophilic MOB, from a boreal lake in Finland. Based on phylogenomic comparisons to MAGs, *Methylobacter* sp. S3L5C represented a ubiquitous cluster of *Methylobacter* spp. in boreal aquatic ecosystems. At optimal temperatures (3–12 °C) and pH (6.8–8.3), the specific growth rates (μ) and CH_4_ utilization rate were in the range of 0.018–0.022 h^-1^ and 0.66–1.52 mmol l^-1^ d^-1^, respectively. In batch cultivation, the isolate could produce organic acids and the concentrations were elevated after replenishing CH_4_ and air into headspace. The highest concentrations of 4.1 mM acetate, 0.02 mM malate and 0.07 mM propionate were observed at the end of the cultivation period under the optimal operational conditions. The results herein highlight the key role of *Methylobacter* spp. in regulating CH_4_ emissions and their potential to provide CH_4_-derived organic carbon compounds to surrounding heterotrophic microorganisms in cold ecosystems.

## 1. Introduction

Methane (CH_4_) is one of major natural and anthropogenic greenhouse gases (GHG), with global warming potential of approximately thirty times higher than that of CO_2_ over a 100-year time horizon (IPCC, 2021). Lakes, one of the major natural sources of CH_4_ emissions, have recently gained interested as they account for ∼6–7% of the total natural emissions (Saunois et al., 2020; Wik et al., 2016). In particular, the emissions from boreal and arctic lakes are elevated during the ice-free period than other times of the year (Guo et al., 2020; Matthews et al., 2020; Sieczko et al., 2020; Wik et al., 2016). Along the lake water column, methanogenic archaea in anoxic sediments/layers produce CH_4_ which is eventually emitted upwards towards the water-atmosphere interface. Various studies have reported the function of aerobic methane oxidizing bacteria (MOB) as the key factor in regulating these CH_4_ fluxes at the oxic-anoxic interfaces in boreal and arctic lake ecosystems (Rissanen et al., 2018, 2021b; Samad and Bertilsson, 2017; Smith et al., 2018; van Grinsven et al., 2021a).

MOB require oxygen as electron acceptor to oxidize CH_4_ for biomass formation and CO_2_ generation. During hypoxic (i.e. oxygen-limiting) conditions, MOB may shift the cellular metabolism towards fermentation to stabilize their cellular redox potential, either by generating various extracellular organic acids (Kalyuzhnaya et al., 2013) or using nitrate as the terminal electron acceptor via anaerobic respiration (Kits et al., 2015). The metabolism of MOB is of great ecological importance as the produced byproducts can serve as growth substrates for surrounding methylotrophic and heterotrophic microorganisms (Ho et al., 2014; Krause et al., 2017). Studies on implementing the produced extracellular byproducts as biofuel-precursors or as industrial platform chemicals also imply the biotechnological importance of MOB (Khanongnuch et al., 2022; Pieja et al., 2017; Strong et al., 2015).

The study of metagenome-assembled genomes (MAG) and experimental observations have revealed that MOB in northern lakes have the genetic potential for denitrification and fermentation resulting in organic acid production (Rissanen et al., 2021b; van Grinsven et al., 2021b). In our previous study on genomic characteristics of boreal lake water column MOB, an attempt to demonstrate organic acid production by MOB in northern lakes was conducted via experimental incubation of natural lake water samples (Rissanen et al., 2021b). However, it was not successful likely due to the low concentrations of organic acids. This suggests that the study on fermentation by lake MOB requires the cultivation of an enrichment or isolate of lake MOB. Up to now, CH_4_ oxidation metabolism in cold conditions, including psychrophilic conditions (0–20 °C), has rarely been studied using enrichment or isolates of MOB (Bertoldo et al., 2003; Trotsenko and Khmelenina, 2005).

The study on lake MOB isolates would benefit in understanding their function, environmental distribution and potential to broaden the biotechnological prospects of MOB. Herein, we focused on isolation, characterization, whole genome assembly, and evaluation of organic acid production of a psychrophilic MOB strain isolated from a boreal lake.

## 2. Material and Methods

### 2.1. Sampling site

Lake water sample from small humic and O_2_-stratified lake, Lake Lovojärvi in southern Finland (61° 04’N, 25° 02’E), was used as the isolation source (Rissanen et al., 2021a). The sampling was conducted at the hypoxic layer (5.75 m depth, temperature 5.2 °C, pH 6.5 and dissolved O_2_ concentration < 0.3 mg l^-1^) (Fig. S1). Lake water samples were pre-filtered through a 50 μm mesh to remove larger plankton. The filtered water was collected in 250 mL glass bottles, containing 20% CH_4_ and 80% air (*v/v*) and stored at 5 °C prior to enrichment.

### 2.2. Enrichment and isolation

In this study, nitrate mineral salts (NMS) medium modified from DSMZ medium 921 with an initial pH of ∼6.80 (Table S1) was used as the liquid growth medium of MOB. Solid medium consisted of the NMS liquid medium supplemented with 1.5% Noble agar (Agar-Agar SERVA powder analytical grade, Germany). For MOB enrichment, 0.5 ml lake water was added into a 25 ml sterile serum bottle containing 5 ml sterile NMS medium. The bottle was sealed with sterile butyl rubber stoppers and aluminum crimps, and the headspace was replaced with 20% (*v/v*) of CH_4_ prior to incubating in static conditions at 5 °C for 42 days. The enriched culture was sub-cultured thrice in NMS medium prior to streaking onto NMS agar medium. Agar plates were incubated in air-tight chamber containing ∼20% CH_4_ and 80% air (*v/v*) in headspace and placed at 5 °C for ∼60 days. Colonies were picked under a stereo microscope and re-streaked on to NMS agar medium. Heterotroph contamination was checked by streaking colonies on nutrient-rich agar medium (0.5% tryptone, 0.25% yeast extract, 0.1% glucose and 2% agar). Culture purity was confirmed with the observation of one cell type under a light microscope and an absence of growth on both nutrient-rich medium and NMS medium without CH_4_ supplementation. The culture purity was also confirmed by high-throughput full-length 16S rRNA gene amplicon sequencing using the Pacbio Sequel platform (Novogene Co. Ltd., Beijing, China).

### 2.3. Physiological tests

The growth tests at different pH, temperatures and nitrogen sources were conducted in 27 mL sterile glass tubes containing 5 mL NMS medium and an initial culture of optical density at 600 nm wavelength (OD_600nm_) of 0.02. The tubes were subsequently sealed with sterile butyl rubber stoppers and aluminum crimps. The initial headspace gas composition of 20% CH_4_ and 80% air (*v/v*) was established by replacing 20% (*v/v*) of the air headspace with pure CH_4_. The pH test was performed in triplicates (pH 4.7–8.3) at 5 °C under static conditions. The growth temperature test was conducted in duplicate using a temperature-gradient incubator (Terratec Corporation, Hobart, Australia), set in the range of 0–26 °C (at pH 6.8). In the different nitrogen source test, NMS and ammonium mineral salts (AMS) media (Table S1) were used as nitrate and ammonium sources, respectively, to cultivate the isolate. The test was conducted in duplicates. Additionally, the total cellular carbohydrate content during the growth in NMS and AMS media was measured at the end of the test. All tests were incubated at static conditions for ∼19–21 days. CH_4_ and CO_2_ compositions in headspace, and OD_600nm_ were monitored daily during weekdays, while O_2_ in the headspace and organic acid concentrations in liquid medium were measured at the end of the test. The growth rates (μ) were obtained from linear regression of the plot between Log_10_ optical density versus incubation time. To test nitrogen (N_2_) fixation, the isolate was incubated in 25 mL serum bottles containing sterile nitrate free–NMS liquid medium. The headspace (*v/v*) was supplemented with (i) 20% CH_4_ + 80% N_2_ and (ii) 20% CH_4_ + 5% O_2_ + 75% N_2_ for anaerobic and microaerobic conditions, respectively. The bottles were sealed with sterile butyl rubber stoppers with aluminum crimps and incubated statically at ∼5 °C for over 30 days.

### 2.4. DNA Extraction and identification

Genomic DNA (gDNA) extraction was performed using GeneJET genomic DNA purification kit (Thermo Fisher Scientific, USA). The gDNA was quantified using Qubit 3.0 Fluorometer and a dsDNA HS Assay Kit (Thermo Fisher Scientific, USA). PCR and amplicon sequencing (Sanger Sequencing method) of the 16S rRNA gene were performed using the identification service offered by Macrogen (Amsterdam, The Netherlands), using primer pairs 27F (AGAGTTTGATCMTGGCTCAG) and 1492R (TACGGYTACCTTGTTACGACTT) for amplification and primer pairs 785F (GGATTAGATACCCTGGTA) and 907R (CCGTCAATTCMTTTRAGTTT) for sequencing. A 16S rRNA gene based phylogenetic tree was constructed in Mega X using the maximum likelihood algorithm (generalized time reversible model) with 100 bootstraps (Kumar et al., 2018). Besides reference strains, the tree was supplemented with 16S rRNA gene sequences of previously studied environmental MAGs representing *Methylobacter* spp. of lakes and ponds of boreal, subarctic and temperate areas (Buck et al., 2021; van Grinsven et al., 2020). For dataset on multiple lakes and ponds, we chose the representative MAGs of metagenomic Operational Taxonomic Units [mOTUs, classified in Buck et al. (2021) at 95% average nucleotide identity cutoff]. The 16S rRNA genes were extracted from MAGs using barrnap (v 0.9) (Seemann, 2018).

### 2.5. Genome sequencing, assembly, and annotation

Library preparation and sequencing for long reads (PacBio Sequel SMRT Cell 1M v2) and short reads (Illumina NovaSeq 6000 platform) were done by Novogene Co. Ltd. (Beijing, China) as previously described by (Rissanen et al., 2021a). The genomes were *de novo* assembled using hybrid assembly strategy in Unicycler (version 0.4.8) with default parameters and “–mode normal” (Wick et al., 2017). The assembled genome was annotated using Prokka (version 1.13.3) (Seemann, 2014). Key functional genes and metabolic pathways of the annotated genome were also predicted and reconstructed using Kyoto Encyclopedia of Genes and Genomes (KEGG) mapping tools (Kanehisa et al., 2022) with KofamKOALA (https://www.genome.jp/tools/kofamkoala/; accessed on 1 February 2022) (Aramaki et al., 2020).

The genome-wide phylogenetic tree was built from protein alignments generated in PhyloPhlAn (v.3.0.58; PhyloPhlAn database including 400 universal marker genes, and “-diversity low” - argument) (Asnicar et al. 2020) using the maximum-likelihood algorithm (PROTCATLG−model) with 100 bootstrap replicates in RAxML (v.8.2.12) (Stamatakis, 2014). Similar to 16S rRNA gene – based phylogenetic tree analysis as explained above, this analysis was supplemented with environmental MAGs representing *Methylobacter* spp. from lakes and ponds of boreal, subarctic, and temperate areas (Buck et al., 2021; van Grinsven et al., 2020), as well as from temperate wetland (Smith et al., 2018). The MAGs were also functionally annotated using Prokka and KofamKOALA as explained above for the genome annotation (Aramaki et al., 2020; Seemann, 2014). Average nucleotide identities (ANI) of the genome with the reference genomes and MAGs were computed using fastANI (version 1.32) (Jain et al., 2018), while average amino-acid identity (AAI) were calculated using CompareM v0.1.2 (https://github.com/dparks1134/CompareM). Digital DNA-DNA hybridization (dDDH) for genome-based taxonomic classification was calculated using the Type Strain Genome Server (TYGS) online service (https://tygs.dsmz.de/; accessed on 28 February 2022) (Meier-Kolthoff and Göker, 2019). Furthermore, *pmoA* sequences, coding for beta subunit of particulate methane monooxygenase, of the MAGs, genome and reference genomes were subjected to phylogenetic tree analyses using the neighbor joining method (Jones-Taylor-Thornton model) with 500 bootstraps in Mega X (Kumar et al., 2018).

### 2.6. Evaluation of organic acid production by the isolate

The organic acid production potential of the isolate was evaluated in batch in 120 mL sterile serum bottles. Prior to sealing with sterile butyl rubber stoppers and aluminum crimps, 15 mL of NMS medium (pH 6.8) were added to the bottles. The initial culture inoculated into the medium and CH_4_ concentration in headspace were previously described in the section 2.5. The experiment was conducted in 6 bottles and incubated at 10.0 ± 1.0 °C in static conditions for 20 days. On day 20, one set of the experiment (in triplicates) was replenished with 20% CH_4_ + 80% air (*v/v*) in headspace, whereas another set (in triplicate) was used as a control without replenishment. The incubation was subsequently continued at the similar conditions until day 33. In this study, organic acid production, OD_600nm_, pH in liquid medium and the gas composition in headspace were periodically monitored every 2 or 3 days. However, in the control test without the replenishment, the gas composition in headspace was monitored only at the beginning and end of the experiment. NMS medium without cells and an inoculated culture without CH_4_ and air addition were also used as negative controls.

### 2.7. Analytical methods

Cell growth was determined using an Ultrospec 500 pro spectrophotometer (Amersham Biosciences, UK). The medium pH was measured using a pH 330i portable meter (WTW, Germany) equipped with a SlimTrode electrode (Hamilton, Germany). To determine the organic acid composition, the cultures centrifuged (15 min at 2700×g) and filtered through a 0.2 μm membrane (Chromafil® Xtra PET 20/25, Macherey-Nagel, Germany) were analyzed using a Shimadzu high-performance liquid chromatograph (HPLC) equipped with Rezex RHM-Monosaccharide H^+^ column (Phenomenex, USA) as described in Okonkwo et al. (2018). Gas composition in headspace (i.e., CH_4_, CO_2_ and O_2_) was measured using a Shimadzu gas chromatograph GC-2014 equipped with a thermal conductivity detector (TCD) and a Carboxen-1000 60/80 column (Agilent Technologies, USA) as described in Khanongnuch et al. (2022). The total carbohydrate content was evaluated using phenol-sulfuric acid method (Masuko et al., 2005). The standard curve of the test was prepared using known glucose concentrations.

### 2.8. Statistical analysis

In each test, the specific growth rate, CH_4_ utilizing rate and organic acid concentration from different experimental conditions were statistically compared using one-way of variance ANOVA (Minitab16.0, USA) with Tukey’s multiple comparison tests. The significance level was the 95% confidence interval, where *p-value* ≤ 0.05 was considered statistically significant.

### 2.9. Accession number

The 16S rRNA gene sequence of *Methylobacter* sp. S3L5C is deposited at GenBank under accession number OM479427. The draft genome assembly of *Methylobacter* sp. S3L5C is available at GenBank under accession number CP076024. The raw reads of the genome sequence are deposited in Sequence Read Archive (SRA) data under accession SRR14663858 for short reads and SRR14663859 for long reads.

## 3. Results

### 3.1. Isolation, characterization, and classification

The isolate was obtained from a single colony forming on NMS agar media statically incubated in an airtight chamber, with ∼20% CH_4_ and 80% air (*v/v*) in the headspace, at ∼5 °C for ∼40 days. The isolate colonies were tiny, less than 0.1 mm diameter, cream, round and entire (Fig. S2a). The cells were small and non-motile cocci (1.0–1.8 μm in diameter) which reproduced by binary fission (Fig. S2b).

For 16S rRNA gene-based identification, the isolate strain S3L5C showed 99.5% and 98.9% similarity to *Methylobacter psychrophilus* Z-0021 and *Methylobacter tundripaludum* SV96 isolated from tundra soil (Omelchenko et al., 1996) and arctic wetland soil (Wartiainen et al., 2006), respectively (Fig. 1; Table S2). In the 16S rRNA gene tree, the strain formed a separate cluster together with *M. psychrophilus* and MAGs representing MOB from lakes and permafrost thaw ponds (Fig. 1). The isolate strain S3L5C was classified in the class *Gammaproteobacteria*, order *Methylococcales*, the family *Methylococcaceae*, and genus *Methylobacter*. Based on the *pmoA* gene-based analysis, the isolate was closely clustered with *M. psychrophilus* and clustered with MAG representing MOB in boreal lake ecosystems (Fig. S3).

**Fig. 1.**
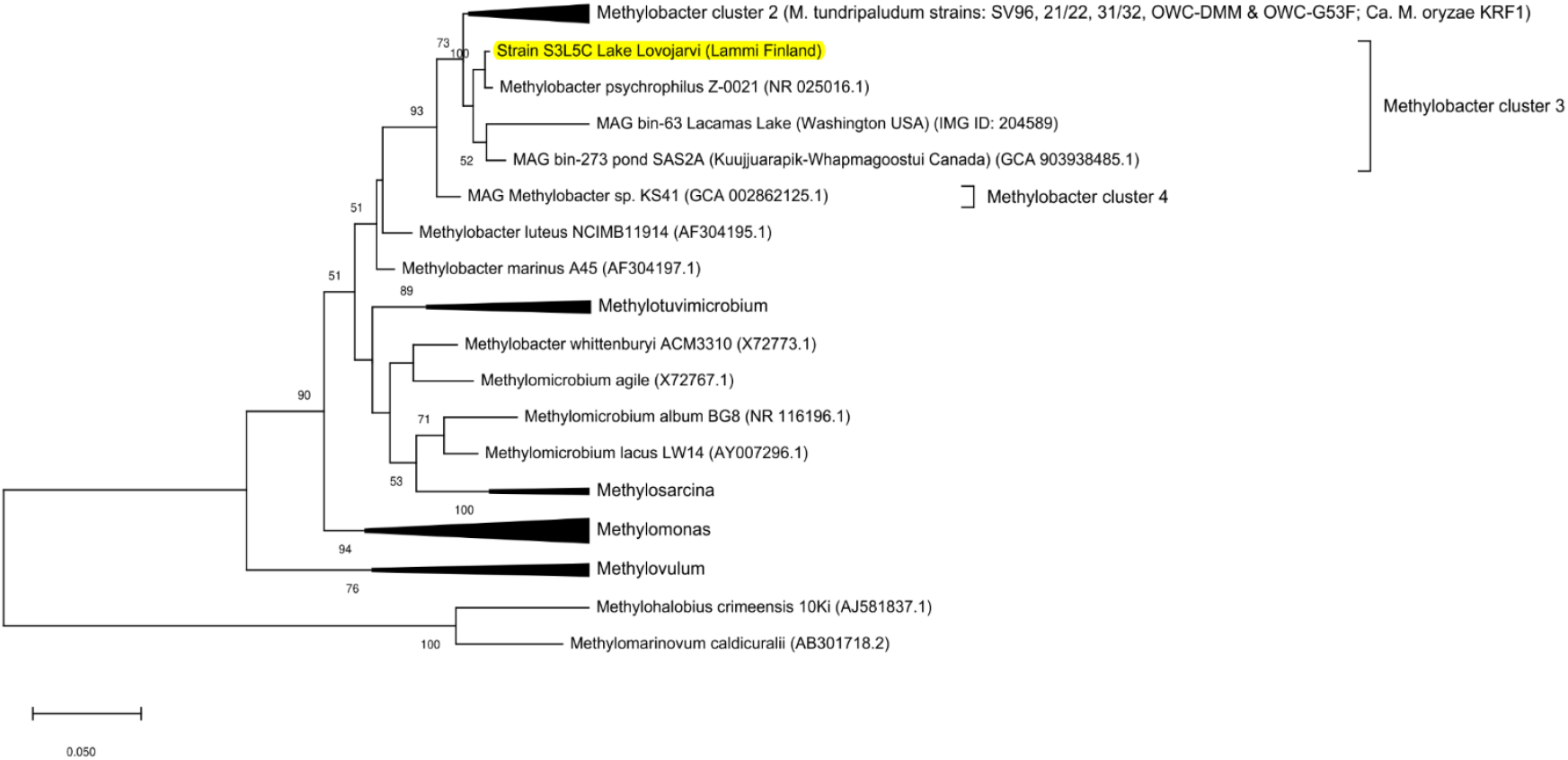
Phylogenetic tree based on 16S rRNA genes strain S3L5C (highlighted in yellow) in comparison with other pure culture methanotrophic bacteria and metagenome-assembled genomes (MAG). GenBank accession numbers are given in parentheses and bar shows 5% sequence divergence.

### 3.2. Physiology

Table 1 shows characteristics of *Methylobacter* sp. S3L5C compared to other psychrophilic and psychrotolerant methanotrophic species. During the cultivation, *Methylobacter* sp. S3L5C grew only in the presence of CH_4_ and O_2_. The growth was not observed in the N_2_-fixation test and under anaerobic conditions, but the cells were still intact in those conditions, even after over 1 year incubation. The strain grew well at 3–12 °C but the cell growth was not observed at above 20 °C, demonstrating psychrophilic nature (Fig. 2a, b). *Methylobacter* sp. S3L5C could grow well in the pH range of 6.0–8.3 and cell growth was not observed at pH 4.6 (Fig. 2d). At the optimal temperatures (3–12 °C) and pH (6.8–8.3), the specific growth rates (μ) and CH_4_ utilization rate were in the range of 0.018–0.022 h^-1^ and 0.66–1.52 mmol l^-1^ d^-1^, respectively (Fig. 2a, b, d, e). Formate (0.04–0.30 mM) and acetate (0.01–0.24 mM) were identified as the major liquid metabolites in all the tested conditions where the strain growth was present (Fig. 2c, f). The trend of acetate production showed that the increased concentration corresponded with the increase of cell growth (Fig. 2c, f).

**Table 1.** Characteristics of different psychrophilic and psychrotolerant species of methanotrophs.

| Strain | *Methylobacter*<br>sp. S3L5C | *Methylobacter*<br>*psychrophilus*<br>Z-0021 | *Methylobacter*<br>*tundripaludum*<br>SV96 | *Methylosphaera*<br>*hansonii* |
| --- | --- | --- | --- | --- |
| Source/habitat | Lake water<br>layer | Tundra soil | Arctic wetland<br>soil | Sediments of<br>marine-salinity<br>Antarctic<br>meromictic<br>lakes |
| Cell morphology | Cocci | Cocci | Rods | Cocci |
| Cell size (µm) | 1.0–1.8 | 1.0–1.7 | 0.8–1.3 × 1.9–<br>2.5 | 1.5–2.0 |
| Motility | - | - | - | - |
| Pigmentation | - | - | Pale Pink | NA |
| Growth pH (optimal) | 6.0–8.3<br>(6.0–7.3) | 5.9–7.0<br>(6.7) | 5.5–7.9 | NA |
| Growth temp. (optimal) (°C) | 0.1–20 (8–12) | 1–20 (10) | 5–30 (23) | 0–21 (10–13) |
| Specific conditions for<br>growth | - | - | - | required sea<br>water |
| Nitrogen fixation gene (nif) | + | NA | + | + |
| G+C content (mol %) | 43.3 | 45–46<sup>†</sup> | 47 | 43–46<sup>†</sup> |
| Reference | This study | (Omelchenko<br>et al., 1996) | (Wartiainen et<br>al., 2006) | (Bowman et al.,<br>1997) |

Note: <sup>†</sup>G+C content was chemically determined as its genome is not available; NA, not available.

**Fig. 2.**
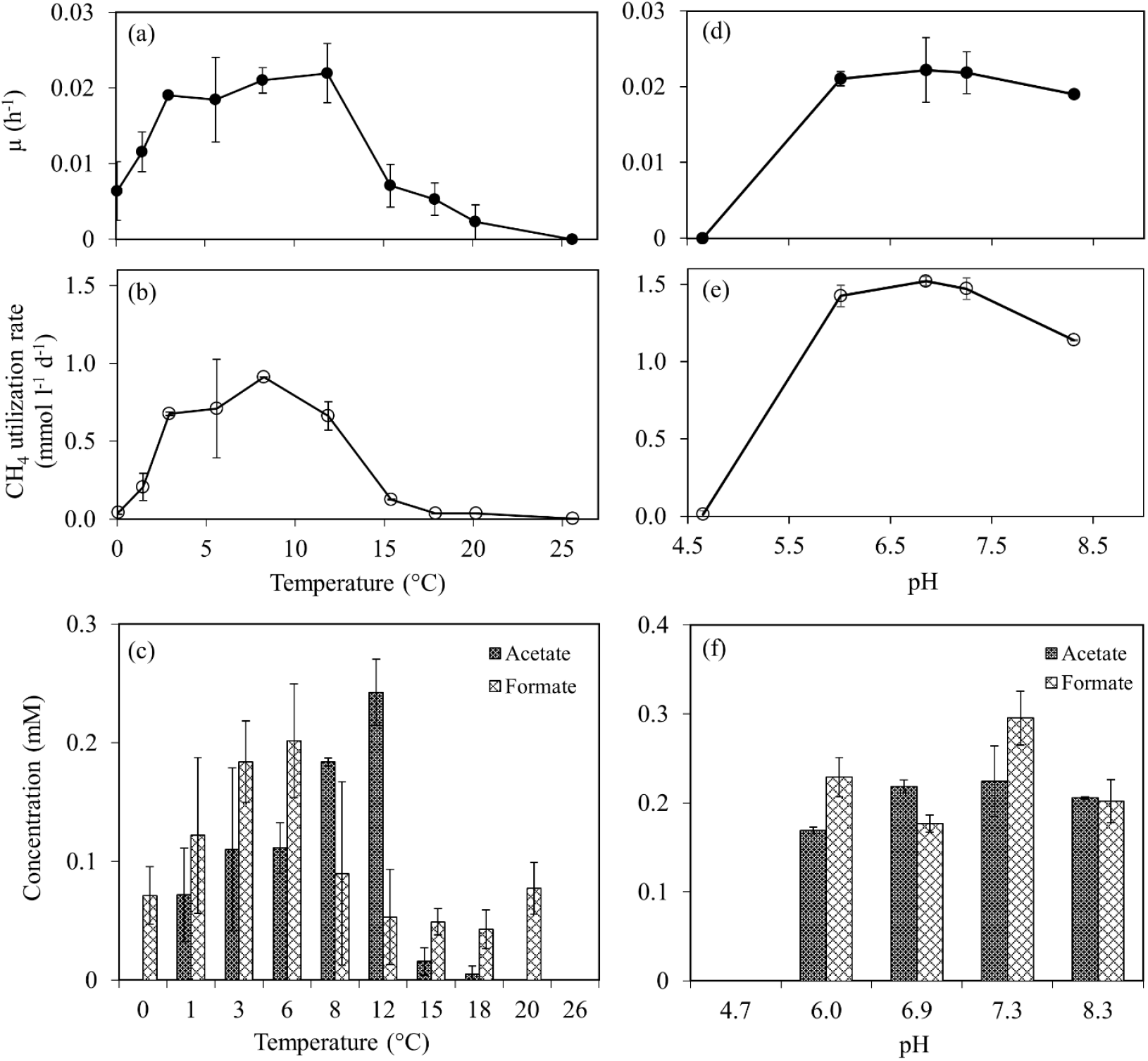
Specific growth (a, d), CH_4_ utilization rate (b, c) and organic acid concentrations (c, f) at the end of the tests at different temperatures (a–c) and pH (d–f). Error bars indicate the standard deviation of duplicates and triplicates for temperature and pH tests, respectively.

*Methylobacter* sp. S3L5C could utilize both nitrate and ammonium as the nitrogen source. The specific growth rate in AMS medium (μ = 0.040 h^-1^) was higher than in NMS medium (μ = 0.020 h^-1^), whereas CH_4_ utilization rate was similar in both NMS and AMS media (0.52–0.88 mmol l^-1^ d^-1^) (Fig. 3a, b). In case of liquid metabolites, acetate and formate were present in NMS medium, while only acetate was detected from the cultivations in AMS medium (Fig. 3c). Furthermore, the total carbohydrate content in biomass cultivated in NMS medium was 0.91 ± 0.04 mM of glucose which was significantly higher than in AMS medium (0.31 ± 0.09 mM of glucose) (*p-value* < 0.001) (Fig. 3d).

**Fig. 3.**
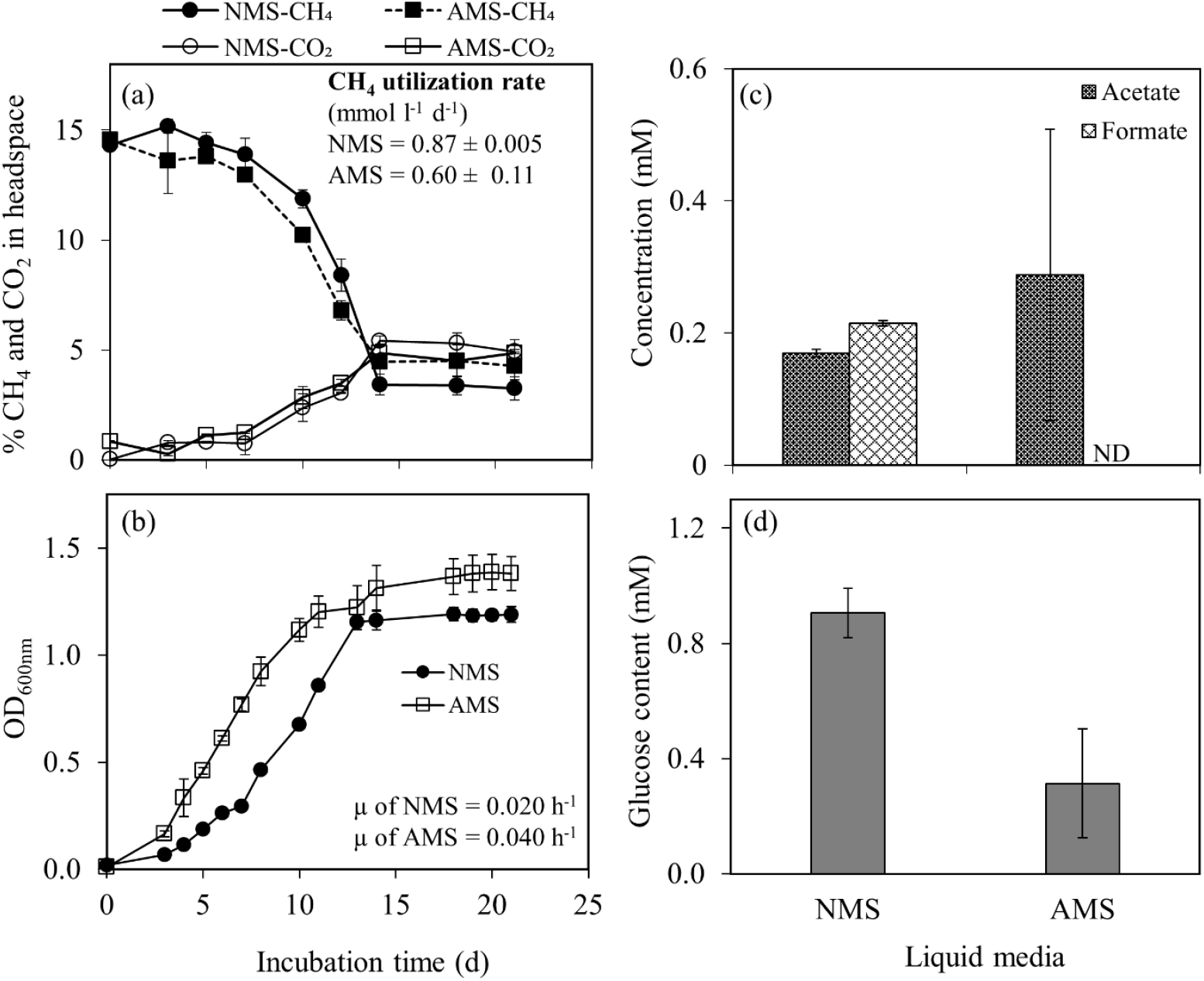
Profiles of (a) CH_4_ utilization and CO_2_ production and (b) biomass growth during 21-day incubation in nitrate (NMS) and ammonium (AMS) mineral salt media. (c) Organic acid production and (d) glucose (representing carbohydrate) content of biomass. Error bars indicate the standard deviation of duplicate samples.

### 3.3. Genome features of *Methylobacter* sp. S3L5C

*Methylobacter* sp. S3L5C genome contained a single chromosome of 4,815,745 bp (G + C content, 43.3%) consisting of 4342 protein-coding sequences, five copies of rRNA (5S, 16S, 23S rRNA), 49 tRNA genes and 2 CRISPR sequences. *Methylobacter* sp. S3L5C genome was clustered together with MAGs of MOB from lakes and ponds in Finland, Sweden, USA, Switzerland and Canada (Fig. 4). Compared to available genomes of other methanotrophic isolates, the S3L5C genome formed a separate species-level branch, and it was not represented by the other isolates (Fig. 4). Regarding similarity indexes for genomic comparison between the SL35C and other MOB, the dDDH was < 25%, while the ANI and AAI values were < 85% (Table S3). To recognize the genomic uniqueness of a novel species, same-species delineation thresholds should be below 70% for dDDH, 95% for ANI, 85% for AAI and 98.7% similarity with 16S rRNA gene sequences (Goris et al., 2007; Khatri et al., 2021; Kim et al., 2014; Luo et al., 2014; Orata et al., 2018; Stackebrandt and Ebers, 2006). Albeit the high sequence similarity of 16S rRNA genes between *Methylobacter* sp. S3L5C and *M. psychrophilus* Z-0021, we could not confirm whether the strains represented identical species due to the non-existence of genome data of *M. psychrophilus* Z-0021. The latter strain is neither available in DSMZ repository nor in All-Russia Collection of Microorganisms (VKM B-2103) culture collection resource, where it was originally deposited (Omelchenko et al., 1996; Wartiainen et al., 2006).

**Fig. 4.**
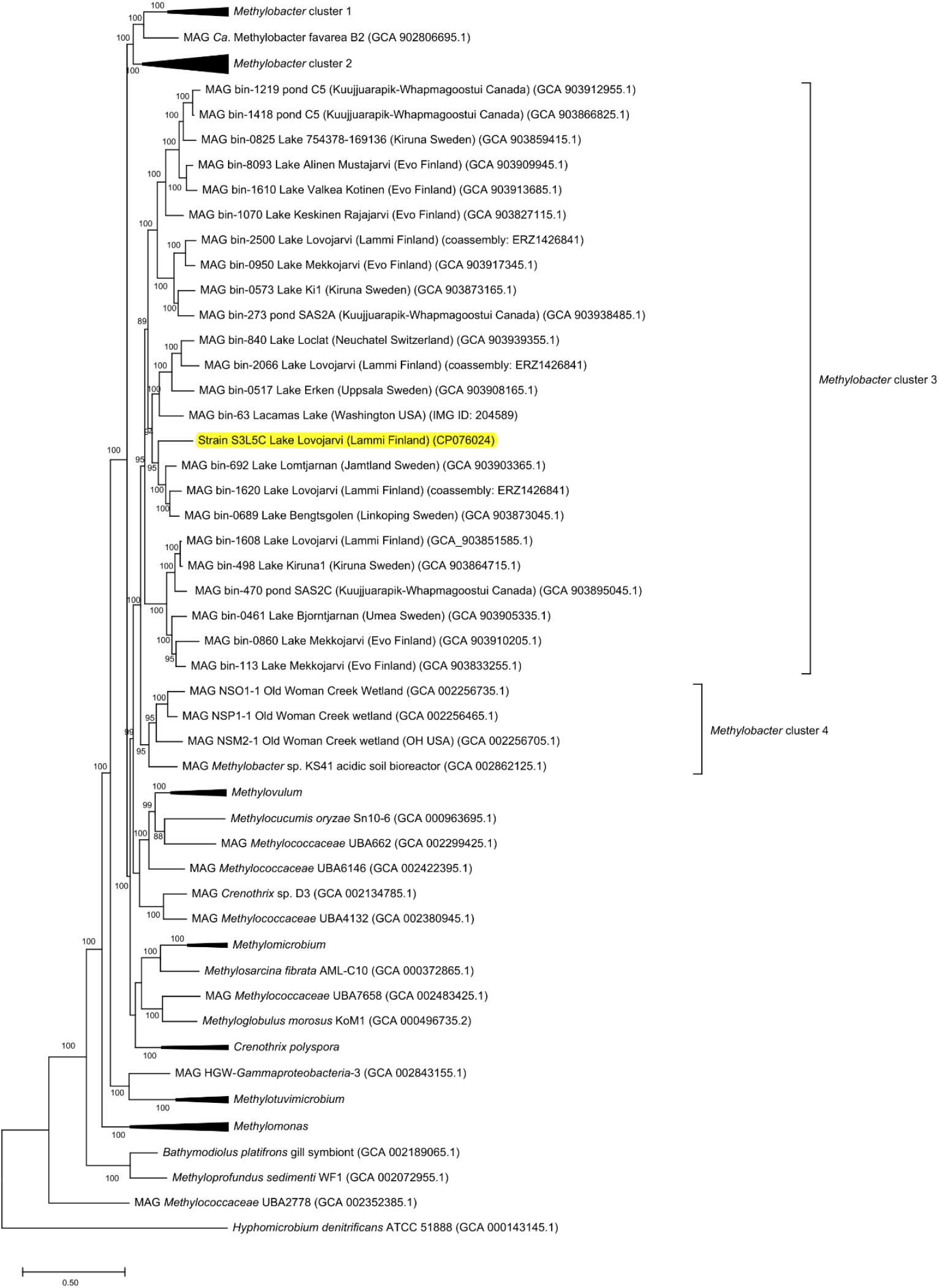
Genome-wide phylogenomic tree constructed using PhyloPhlAn2 showing the position of *Methylobacter* sp. S3L5C (highlighted in yellow) compared to other cultured methanotrophs and metagenome-assembled genomes (MAGs) of uncultured *Methylobacter* and *Methylococcales* from environment samples. *Methylobacter* clusters 1 and 2 contain the so far cultured members of *Methylobacter* spp.

### 3.4. Predicted metabolic pathways

The key metabolic pathways in *Methylobacter* sp. S3L5C were predicted based on the KEGG database (Fig. 5). *Methylobacter* sp. S3L5C genome contains all key genes associated with CH_4_ oxidation including both particulate (*pmoCAB*) and soluble (*mmoXYBZDC*) methane monooxygenases. For the conversion of methanol to formaldehyde, the strain contained both calcium- (*mxaFJGIACKLD*) and lanthanide-dependent (*xoxF*) methanol dehydrogenases. Genes encoding tetrahydromethanopterin (H_4_MPT)-mediated pathway, catalyzing the conversion of formaldehyde into formate, were present in the isolate. *Methylobacter* sp. S3L5C contained a complete set of genes encoding major pathways for formaldehyde assimilation into cell biomass including ribulose monophosphate (RuMP), Embden–Meyerhof–Parnas (EMP), Enter-Doudoroff (EDD) and phosphoketolases (*xfp*)-based pathway (Fig. 5; Table S4). The strain lacked the gene encoding serine-glyoxylate aminotransferase (*sga*) incompleting the serine pathway. Genes present in *Methylobacter* sp. S3L5C also encoded the oxidative tricarboxylic acid (TCA) cycle and C5-branched dibasic acid metabolism.

**Fig. 5.**
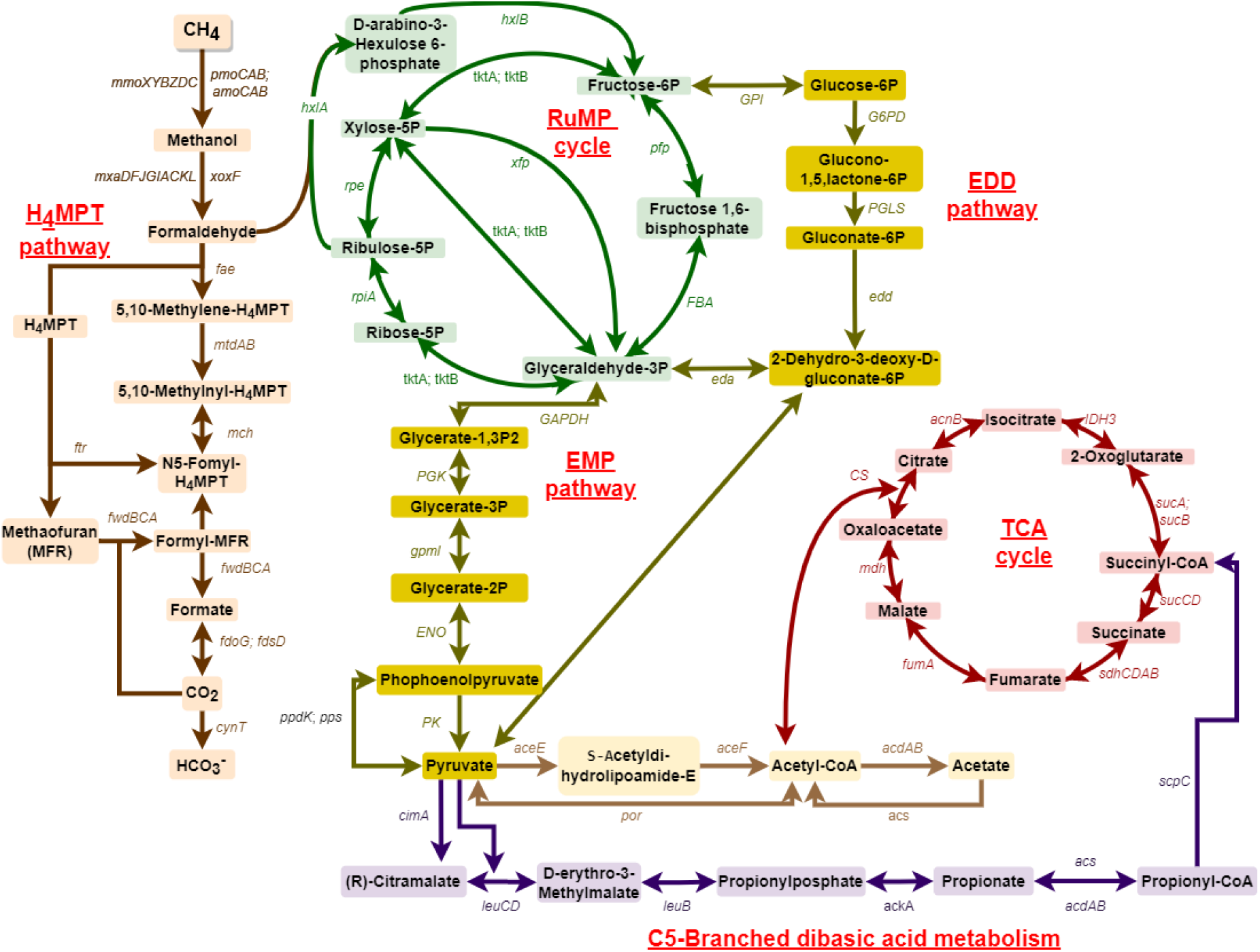
Predicted metabolic pathway of *Methylobacter* sp. S3L5C constructed based on KEGG mapper (Kanehisa et al., 2022). List of genes are present in Table S4. H_4_MPT, tetrahydromethanopterin; RuMP, ribulose monophosphate; EMP, Embden-Meyerhof-Parnas; EDD, Enter-Doudoroff; TCA, tricarboxylic acid.

Gammaproteobacterial MOB generally can oxidize NH_4_^+^ into hydroxylamine (NH_2_OH) using pmoCAB (Mohammadi et al., 2017). Our experimental observation suggests that the same NH_4_^+^ metabolism might also occur in *Methylobacter* sp. S3L5C. However, the latter did not contain putative *hao* genes encoding hydroxylamine oxidoreductase to convert hydroxylamine to nitrite or nitric oxide (Versantvoort et al., 2020). The genome also included genes related to nitrate/nitrite transporter i.e., assimilatory nitrate reductase (*nasA*) for reducing nitrate into nitrite, nitrite reductase (*nirBD*, and *nirS*) for reducing nitrite into ammonia and nitric oxide, as well as nitric oxide reductase (*norB*) for reducing nitric oxide into nitrous oxide. Genes encoding nitrogen fixation (*nifHDK*) were found, although the CH_4_ oxidation was not observed in nitrogen-free medium tests under anaerobic and microaerobic (5% O_2_) conditions. Genes encoding the assimilatory sulfate reduction (sulfate reduced to sulfide), sulfate/sulfur reductase (*cysJNCDIH*), were also present in *Methylobacter* sp. S3L5C.

Genes encoding the key enzymes involved in fermentative metabolisms were observed in the genome, including acetyl-CoA synthetase (*acdAB*) catalyzing the conversion of acetyl/propionyl–CoA into acetate/propionate and acetate kinase (*ackA*) for propionate generation from propionyl phosphate. However, some genes encoding acetate formation were not present in the genome, such as phosphate acetyltransferase (*pta*) and pyruvate dehydrogenase (*poxB*). The genome also included malate dehydrogenase (*mdh*) encoding the reversible conversion of oxaloacetate, succinate dehydrogenase (*sdhCDAB*) coupling with succinate production and NAD^+^-reducing hydrogenase (*hoxFGYH*) encoding H_2_ production. Based on further analyses of environmental MAGs, similar genetic potential for fermentation and organic acid production is widely dispersed in closely related *Methylobacter* spp. of lakes, ponds, and wetlands of temperate and boreal area (Table S5).

### 3.5. Organic acid production

Batch cultivations were performed to evaluate the strain’s potential to produce organic acids by replenishing CH_4_ and air during incubation (on day 20, Fig. 6a). In the CH_4_+air replenishment test, the average utilized oxygen-to-methane (O_2_/CH_4_) molar ratio was 1.0 ± 0.5 mol mol^-1^ during cultivation from days 4–33 (Fig. S4). At the end of the test (day 33), the culture with the replenished headspace had utilized CH_4_ and O_2_ concentrations at ∼1.5–2 times higher than those of the control cultivations without the replenishment, corresponding with the higher CH_4_ and O_2_ concentrations fed into the system (Fig. S5a). On day 33, the CH_4_+air replenishment also increased the biomass growth (up to OD_600nm_ 5.0 ± 0.2), while an OD_600nm_ of 3.2 ± 0.4 was obtained in the control cultivations without CH_4_+air replenishment (Fig. S5b). Although similar CH_4_ utilization efficiency (64.1–69.3%) were observed, the control demonstrated higher O_2_ utilization efficiency (72%) that the test with CH_4_+air replenishment (*p-value* = 0.041) (Table S6).

**Fig. 6.**
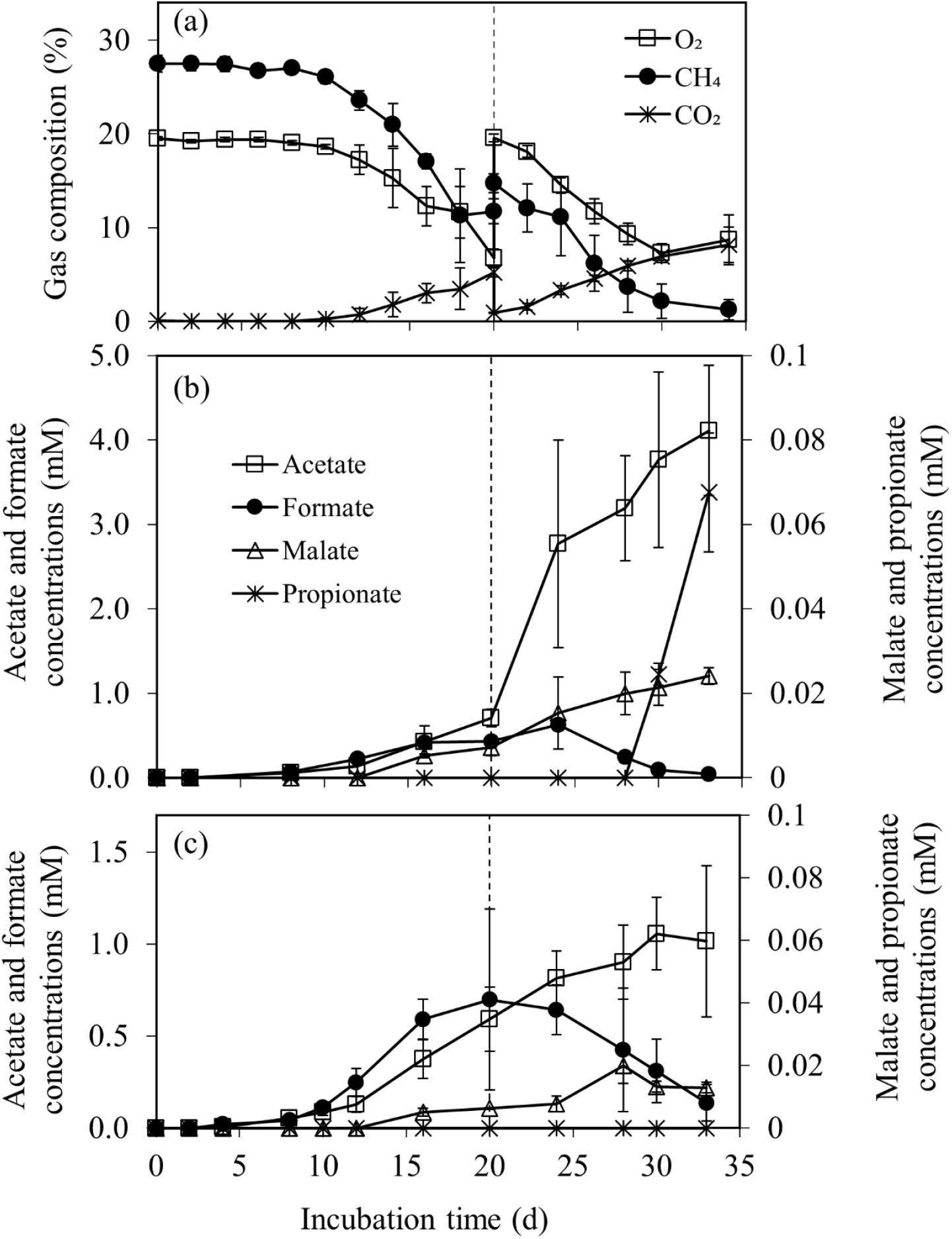
The profiles of (a) gas composition in headspace of the test with 20% CH_4_ and 80% air replenishment on day 20 and concentrations of organic acids in liquid medium during 33-day cultivation of strain S3L5C (b) with and (c) without the replenishment. Error bars indicate the standard deviation of triplicate samples.

Acetate, formate, and malate were observed as soluble metabolites from *Methylobacter* sp. S3L5C cultivations with and without the replenishment, while only propionate was present in the test with CH_4_ and air replenishment (Fig. 6b). Organic acid concentrations gradually accumulated during the cultivation period, except for formate which was mostly depleted in both experimental conditions (Fig. 6b, c). In the tests with CH_4_+air replenishment, the accumulated concentrations of acetate (4.1 mM), malate (0.02 mM) and propionate (0.07 mM) were significantly higher than those in the control (*p-value* < 0.05) (Fig. 6b, c). In this study, *Methylobacter* sp. S3L5C is hypothesized to convert carbon from CH_4_ (C-CH_4_) into organic acids, CO_2,_ and biomass. The carbon conversion into total accumulated organic acids in the test with CH_4_+air replenishment (2.5% of the consumed C-CH_4_) was significantly higher than those in the control (1.2% of the consumed C-CH_4_) (*p-value* = 0.037) (Table S7). In both experimental conditions, the carbon yields of C-CO_2_ and C-biomass derived from C-CH_4_ were similar in the range of 42.2–46.3% and 51.1–56.4%, respectively (*p-value* = 0.124) (Table. S7).

## 4. Discussion

Strain S3L5C, isolated and classified in this study as *Methylobacter* sp., has been previously reported to be a dominant genus at oxic-anoxic transition zone in boreal and subarctic lakes, ponds and wetlands (Cabrol et al., 2020; Cassarini et al., 2019; Martin et al., 2021; Rissanen et al., 2021b, 2018; Smith et al., 2018; van Grinsven et al., 2021a). Our phylogenetic and phylogenomic tree analyses showed that the isolate represents a ubiquitous cluster of *Methylobacter* spp. in lake and pond ecosystems. *Methylobacter* sp. S3L5C genome contained genes encoding key enzymes involved in aerobic CH_4_ oxidation and organic acid production (fermentative metabolism). According to previous studies and MAG analyses of this study, genetic potential of *Methylobacter* spp. for organic acid production is widely dispersed in CH_4_-rich aquatic systems (Table S5) (Rissanen et al., 2021b; Smith et al., 2018; van Grinsven et al., 2020). Whether or not *Methylobacter* sp. S3L5C is a new species of *Methylobacter* or a type strain of the previously described *Methylobacter psychrophilus* Z-0021 could not be validated as the strain Z-0021 and its genome is not available. Nevertheless, *Methylobacter* sp. S3L5C represents psychrophilic methanotrophs that rarely exist as isolates.

Organic acid production in MOB is an energy conservation mechanism, where pyruvate is oxidized to fermentative by-products instead towards biomass formation during substrate limiting conditions (e.g. CH_4_ and O_2_) (Kalyuzhnaya et al., 2013; Lee et al., 2016). In our study, the average O_2_/CH_4_ uptake ratio during incubation from days 4–33 (O_2_/CH_4_ uptake ratio of 1.0 ± 0.6; Fig. S5) was lower than the stoichiometric ratio in aerobic CH_4_ oxidation (Eq. 1).

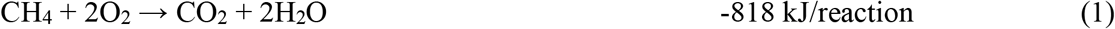

These conditions probably induced CH_4_ oxidation with O_2_ limited amount in the system and initiated the accumulation of acetate, propionate and malate. The O_2_/CH_4_ uptake behavior was similar to the previous studies conducted using *Methylomicrobium buryatense* 5GB1C strain cultivated under O_2_ and CH_4_ limitation conditions (O_2_/CH_4_ uptake ratio of 1.1–1.6) (Gilman et al., 2017, 2015).

When *Methylobacter* sp. S3L5C grew in batch cultivation, acetate was observed as the dominant metabolite and its concentration was remarkably elevated under the conditions with CH_4_+air replenishment in headspace. *Methylobacter* sp. S3L5C genome lacks the genes encoding phosphate acetyltransferase (*pta*) and pyruvate dehydrogenase (*poxB*). Nevertheless, experimental validation on the metabolite production suggests that acetate synthesis route might be catalyzed by acetyl-CoA synthetase and ligase from acetyl-CoA found in the genome (*acs* and *acdAB*; Fig. 5 and Table S4) (Schäfer et al., 1993). To the best of our knowledge, this is the first study to report the capacity of a MOB isolate to produce propionate, which is a liquid metabolite commonly produced by archaea and facultative anaerobic bacteria under anaerobic/microaerobic conditions (Baleeiro et al., 2021; Gonzalez-Garcia et al., 2017; Moreira et al., 2021). The genome of *Methylobacter* sp. S3L5C contained annotations for genes encoding propionate production (*acdAB, acs* and *ackA*). Thus, based on the *in silico* data (Fig. 5; Table S4), we hypothesize that the propionate production from *Methylobacter* sp. S3L5C may occur via succinate and citramalate pathways similar to other propionate producing microorganisms (Gonzalez-Garcia et al., 2017). Formate accumulation in the liquid culture has been observed during an unbalanced growth conditions (e.g. under O_2_ limitation and cultivated in methanol as a carbon source) (Gilman et al., 2017; Kalyuzhnaya et al., 2013; Tays et al., 2018). However, in our study, formate was observed as an intermediate metabolite which was eventually depleted during the cultivation period (Fig. 6b, c). Although the genes encoding succinate dehydrogenase was annotated in the genome, succinate was not detected in any studied conditions.

While some MOB are capable of nitrogen fixation, ammonium and nitrate are widely nitrogen sources for MOB biomass assimilation (Hanson and Hanson, 1996; Strong et al., 2015). Our results revealed the effect of different nitrogen sources (i.e., nitrate and ammonium) on *Methylobacter* sp. S3L5C growth. The strain favored ammonium for biomass assimilation, resulting in significantly higher growth rate than the nitrate medium (Fig. 3b). The efficient growth in ammonium medium can be attributed to the original lake environment (high ammonium content, DO < 0.3 mg l^-1^, Fig. S1) from which *Methylobacter* sp. S3L5C was isolated. In the previous study on the bacterial community in anoxic brackish groundwater enrichments, *Methylobacter* sp. were observed as a predominant methanotroph in the CH_4_ and ammonium-rich medium, but it was absent in the nitrate supplemented medium (Kutvonen et al., 2015). However, studies on different gamma- and alphaproteobacterial MOB have reported varying responses on the usage of nitrate or ammonium as the optimum nitrogen source (Kits et al., 2015; Tays et al., 2018). For instance, Tays et al. (2018) reported that a medium containing ammonium was optimal for the growth of *Methylocystis sp*. Rockwell, albeit with lower lipid content (fatty acid methyl ester, FAME) compared to the growth in nitrate medium (Tays et al., 2018). In some circumstances, nitrate favored the growth of *Methylomonas denitrificans* FJG1 under O_2_-limited conditions (Kits et al., 2015). In our study, the strain’s biomass collected from the ammonium medium had significantly less carbohydrate (sugar) content than the cells growing on the nitrate medium (Fig. 3d). This result suggests that the use of different nitrogen sources can be adopted as a strategy to target the major biomass composition (protein, lipid, and carbohydrate contents) during *Methylobacter sp*. S3L5C cultivation. In methanotrophic biomass, carbohydrate commonly represents a carbon sink which might not be preferable for biotechnological applications (e.g., single cell protein and microbial lipid derived fuels) (Fei et al., 2018; Gilman et al., 2015; Nguyen and Lee, 2020).

In conclusion, our results on *Methylobacter* sp. S3L5C and comparative MAG analyses suggest that *Methylobacter* spp. play a key role in mitigating of atmospheric CH_4_ emissions and synthesizing organic acids in boreal and subarctic aquatic ecosystems. The findings suggest that the organic acids produced by *Methylobacter* spp. could be served as a carbon source for surrounding heterotrophic microorganisms to sustain a functioning ecosystem (Ho et al., 2014; Kankaala et al., 2006). Furthermore, *Methylobacter* sp. S3L5C may provide an alternative source of CH_4_-derived metabolites for biotechnological applications, especially in cold systems. For example, organic acid rich spent media can be used to cultivate recombinant heterotrophs to generate value-added compounds (Khanongnuch et al., 2022; Lee et al., 2021; Takeuchi and Yoshioka, 2021). However, further studies on the effect of alternative electron acceptors (e.g., nitrogen oxides) on metabolism and organic acid production of *Methylobacter* sp. S3L5C and process optimizations are required to enhance the production of CH_4_-derived products.

## Acknowledgements

This research work was funded by Kone foundation [grant number 201803224 for AJR and RK] and Academy of Finland [grant number 323214 for RM]. The authors thank Anne Grethe Hestnes, The Arctic University of Norway, Tromsø, Norway, for her guidance and support in methanotroph isolation and cultivation. The authors also thank the staff at Lammi Biological Station (Finland) for their support in sampling.

