## Supplementary materials for "Characterization and genome analysis of a psychrophilic methanotroph representing a ubiquitous *Methylobacter* spp. cluster in boreal lake ecosystems"

**Table S1.** Composition of nitrate mineral salts (NMS) and trace elements solution used in this study, modified from DSMZ medium 921.

| Composition | Concentration (mg $L^{-1}$) |
|---|---|
| $KNO_3$ ($NH_4Cl$ for AMS[†]) | 1000 (530) |
| $MgSO_4 \cdot 7H_2O$ | 1000 |
| CaCl2·2H2O | 200 |
| $K_2HPO_4$ | 348.4 |
| $KH_2PO_4$ | 260 |
| Fe(III)-EDTA | 3.8 |
| **Trace elements** | |
| $CuSO_4 \cdot 5H_2O$ | 1 |
| $FeSO_4 \cdot 7H_2O$ | 0.5 |
| $ZnSO_4 \cdot 7H_2O$ | 0.4 |
| $H_3BO_3$ | 0.015 |
| $CoCl_2 \cdot 6H_2O$ | 0.05 |
| EDTA-$Na_2$ | 0.25 |
| $MnCl \cdot 4H_2O$ | 0.02 |
| NiCl2·$6H_2O$ | 0.01 |
| $Na_2MoO_4 \cdot 2H_2O$ | 0.23 |
| $LaCl_3$ | 0.16 |

Note: [†]AMS, ammonia mineral salts medium

**Table S2.** 16S rRNA gene sequences of *Methylobacter* strain S3L5C with query length of 1486 bp compared to other pure culture strains identified using Nucleotide BLAST (blastn), National Center for Biotechnology Information (NCBI [https://www.ncbi.nlm.nih.gov/]).

| | **Query cover (%)** | **Identity (%)** | **Length (bp)** | **Accession number** |
|---|---|---|---|---|
| *Methylobacter psychrophilus* Z-0021 | 99 | 99.46 | 1527 | NR_025016.1 |
| *Methylobacter tundripaludum* SV96 | 98 | 98.91 | 1487 | NR_042107.1 |
| *Methylobacter luteus* NCIMB 11914 | 99 | 96.48 | 1500 | AF304195.1 |
| *Methylobacter marinus* A45 | 99 | 96.34 | 1499 | AF304197.1 |
| *Methylobacter whittenburyi* ACM3310 | 97 | 95.24 | 1470 | X72773.1 |
| *Methylomicrobium lacus* LW14 | 99 | 95.13 | 1499 | AY007296.1 |
| *Methylomicrobium agile* | 97 | 94.34 | 1469 | X72767.1 |
| *Methylomicrobium album* BG8 | 97 | 94.05 | 1447 | NR 116196.1 |
| *Methylovulum miyakonense* HT12 | 98 | 92.10 | 1488 | NR_112920.1 |
| *Methylomarinovum caldicuralii* | 98 | 86.97 | 1466 | AB301718.2 |
| *Methylohalobius crimeensis* 10Ki | 98 | 86.58 | 1462 | AJ581837.1 |

**Table S3.** The digital DNA-DNA hybridization (dDDH), average nucleotide identity (ANI) and average amino acid identity (AAI) values of strain S3L5C compared to closely related reference methanotrophic strains and metagenome-assembled genomes (MAG) of methanotrophs in lakes and wetlands of temperate, boreal and subarctic area.

| **S3L5C compared to** | **dDDH** | **ANI value (%)** | **AAI value (%)** |
|---|---|---|---|
| *Methylomonas methanica* NCIMB 11130 (GCA 001644045.1) | 23.1 | n.a. | n.a. |
| *Methylomonas denitrificans* FJG1T | 22.9 | n.a. | n.a. |
| *Methylobacter tundripaludum* SV96 (GCA 000190755.3) | 21.8 | 77.8 | 76.6 |
| Candidatus *Methylobacter oryzae* KRF1 (GCA 003994235.2) | 20.6 | n.a. | 75.1 |
| *Methylobacter marinus* A45 (GCA_000383855.1) | 20.6 | n.a. | 72.6 |
| *Methylosarcina fibrata* AML-C10 (GCA 000372865.1) | 20.5 | n.a. | 69.9 |
| *Methyloprofundus sedimenti* WF1 (GCA 002072955.1) | 20.4 | n.a. | 64.4 |
| *Methylosarcina lacus* LW14 (GCA 000527095.1) | 20.3 | n.a. | 70.3 |
| *Methylocucumis oryzae* Sn10-6 (GCA_000963695.1) | 20.3 | n.a. | 67.4 |
| *Methylomicrobium alcaliphilum* 20Z (GCA 000968535.1) | 20.2 | n.a. | 67.2 |
| MAG bin-692 Lake Lomtjarnan (Jamtland Sweden) (GCA 903903365.1) | - | 82.1 | 84.7 |
| MAG bin-0689 Lake Bengtsgolen (Linkoping Sweden) (GCA 903873045.1) | - | 81.8 | 84.4 |
| MAG NSO1-1 Old Woman Creek Wetland (GCA 002256735.1) | - | 80.7 | 83.7 |
| MAG *Methylobacter* sp. KS41 acidic soil bioreactor (GCA 002862125.1) | - | 79.3 | 80.3 |
| MAG bin-840 Lake Loclat (Neuchatel Switzerland) (GCA 903939355.1) | - | 78.5 | 79.1 |
| MAG bin-63 Lacamas Lake (Washington USA) (IMG ID: 204589) | - | 78.4 | 80.6 |
| bin-0517 Lake Erken (Uppsala Sweden) (GCA 903908165.1) | - | 78.2 | 78.8 |
| MAG bin-0825 Lake 754378-169136 (Kiruna Sweden) (GCA 903859415.1) | - | 78.0 | 78.5 |
| MAG bin-0461 Lake Bjorntjarnan (Umea Sweden) (GCA 903905335.1) | - | 77.8 | 77.0 |
| MAG bin-1219 pond C5 (Kuujjuarapik-Whapmagoostui Canada) (GCA 903912955.1) | - | 77.8 | 77.9 |
| MAG bin-1070 Lake Keskinen Rajajarvi (Evo Finland) (GCA 903827115.1) | - | 77.7 | 77.4 |
| MAG bin-113 Lake Mekkojarvi (Evo Finland) (GCA 903833255.1) | - | 77.7 | 77.2 |
| MAG bin-498 Lake Kiruna1 (Kiruna Sweden) (GCA 903864715.1) | - | 77.6 | 76.5 |

| | | | |
|---|---|---|---|
| MAG bin-273 pond SAS2A (Kuujjuarapik-Whapmagoostui Canada) (GCA 903938485.1) | - | 77.6 | 75.6 |
| MAG bin-1610 Lake Valkea Kotinen (Evo Finland) (GCA 903913685.1) | - | 77.6 | 78.4 |
| MAG bin-8093 Lake Alinen Mustajarvi (Evo Finland) (GCA 903909945.1) | - | 77.6 | 78.2 |
| MAG bin-1418 pond C5 (Kuujjuarapik-Whapmagoostui Canada) (GCA 903866825.1) | - | 77.5 | 77.6 |
| MAG bin-470 pond SAS2C (Kuujjuarapik-Whapmagoostui Canada) (GCA 903895045.1) | - | 77.5 | 76.7 |
| MAG bin-0950 Lake Mekkojarvi (Evo Finland) (GCA 903917345.1) | - | 77.3 | 75.6 |
| MAG bin-0860 Lake Mekkojarvi (Evo Finland) (GCA 903910205.1) | - | 77.1 | 76.1 |
| MAG *Methylobacter* sp. B2 (GCA 902806695.1) | - | n.a. | 74.1 |
| *Methylobacter luteus* IMV-B-3098 (GCA 000427625.1) | - | n.a. | 72.8 |
| *Methylobacter whittenburyi* UCM-B-3033 (GCA_000745375.1) | - | n.a. | 72.4 |
| *Methylobacter* sp. BBA5 (GCA 000746145.1) | - | n.a. | 72.3 |
| *Methylovulum psychrotolerans* HV10-M2 (GCA 002209385.1) | - | n.a. | 71.4 |
| *Methyloglobulus morosus* KoM1 (GCA 000496735.2) | - | n.a. | 71.5 |
| *Methylomicrobium album* BG8 (GCA 000214275.3) | - | n.a. | 69.2 |
| *Methylomicrobium buryatense* 5GB1C (GCA 005931095.1) | - | n.a. | 67.6 |

Note: n.a., not available because ANI values lower than 76.8% were not present in the results using fastANI

**Table S4.** Functional enzymes in *Methylobacter* sp. S3L5C based on KEGG database

| Locus tag | KEGG ID | Gene | Description | Pathway |
|---|---|---|---|---|
| PIAMOCKP_01318 | K10946 | pmoC | particular methane/ammonia monooxygenase subunit C [EC:1.14.18.3] | $CH_4$ oxidation |
| PIAMOCKP_01319 | K10944 | pmoA | particular methane/ammonia monooxygenase beta subunit [EC:1.14.18.3] | |
| PIAMOCKP_01320 | K10945 | pmoB | particular methane/ammonia monooxygenase alpha subunit B [EC:1.14.18.3] | |
| PIAMOCKP_02349 | K16157 | mmoX | soluble methane monooxygenase subunit X [EC:1.14.13.25] | |
| PIAMOCKP_02350 | K16158 | mmoY | soluble methane monooxygenase subunit Y [EC:1.14.13.25] | |
| PIAMOCKP_02351 | K16160 | mmoB | soluble methane monooxygenase subunit B [EC:1.14.13.25] | |
| PIAMOCKP_02352 | K16159 | mmoZ | soluble methane monooxygenase subunit Z [EC:1.14.13.25] | |
| PIAMOCKP_02353 | K16162 | mmoD | soluble methane monooxygenase subunit D [EC:1.14.13.25] | |
| PIAMOCKP_02354 | K16161 | mmoC | soluble methane monooxygenase subunit C [EC:1.14.13.25] | |
| PIAMOCKP_00118 | K23995 | xoxF | lanthanide-dependent methanol dehydrogenase [EC:1.1.2.10] | Methanol oxidation |
| PIAMOCKP_02370 | K16260 | mxaD | methanol dehydrogenase [EC:1.1.2.7] | |
| PIAMOCKP_02372 | K14028 | mxaF | calcium-dependent methanol dehydrogenase, large subunit [EC:1.1.2.7] | |
| PIAMOCKP_02373 | K16254 | mxaJ | methanol dehydrogenase [EC:1.1.2.7] | |
| PIAMOCKP_02374 | K16255 | mxaG | methanol dehydrogenase (cytochrome c electron acceptor) [EC:1.1.2.7] | |
| PIAMOCKP_02375 | K14029 | mxaI | calcium-dependent methanol dehydrogenase, small subunit [EC:1.1.2.7] | |
| PIAMOCKP_02378 | K16256 | mxaA | methanol dehydrogenase (protein for calcium insertion) [EC:1.1.2.7] | |
| PIAMOCKP_02379 | K16257 | mxaC | methanol dehydrogenase (protein for calcium insertion) [EC:1.1.2.7] | |
| PIAMOCKP_02380 | K16258 | mxaK | methanol dehydrogenase (protein for calcium insertion) [EC:1.1.2.7] | |
| PIAMOCKP_02381 | K16259 | mxaL | methanol dehydrogenase (protein for calcium insertion) [EC:1.1.2.7] | |
| PIAMOCKP_01814 | K10713 | fae | formaldehyde-activating enzyme [EC:4.2.1.147] | Formaldehyde |
| PIAMOCKP_00059 | K00300 | mtdA | Methylene-tetrahydromethanopterin dehydrogenase (NADP+) [EC:1.5.1.-] | |
| PIAMOCKP_04319 | K10714 | mtdB | methylene-tetrahydromethanopterin dehydrogenase [EC:1.5.1.-] | |
| PIAMOCKP_01810 | K01499 | mch | methenyltetrahydromethanopterin cyclohydrolase [EC:3.5.4.27] | |
| PIAMOCKP_01347 | K00672 | ftr | Formylmethanofuran-tetrahydromethanopterin N-formyltransferase [EC:2.3.1.101] | |
| PIAMOCKP_01345 | K00201 | fwdB | formylmethanofuran dehydrogenase subunit B [EC:1.2.7.12] | |
| PIAMOCKP_01348 | K00202 | fwdC | formylmethanofuran dehydrogenase subunit C [EC:1.2.7.12] | |
| PIAMOCKP_01678 | K00200 | fwdA | formylmethanofuran dehydrogenase subunit A [EC:1.2.7.12] | |

| Locus tag | KEGG ID | Gene | Description | Pathway |
|---|---|---|---|---|
| PIAMOCKP_04158 | K00123 | fdoG | formate dehydrogenase major subunit [EC:1.17.1.9] | |
| PIAMOCKP_04161 | K00126 | fdsD | formate dehydrogenase subunit delta [EC:1.17.1.9] | |
| PIAMOCKP_03844 | K01673 | cynT | carbonic anhydrase [EC:4.2.1.1] | |
| PIAMOCKP_01399 | K08093 | hxlA | 3-hexulose-6-phosphate synthase [EC:4.1.2.43] | RUMP cycle |
| PIAMOCKP_01400 | K08094 | hxlB | 6-phospho-3-hexuloisomerase [EC:5.3.1.27] | |
| PIAMOCKP_00638 | K00895 | pfp | diphosphate-dependent phosphofructokinase | |
| PIAMOCKP_01406 | K01624 | fbaA | fructose-bisphosphate aldolase, class II | |
| PIAMOCKP_00835 | K00615 | tktAB (E2.2.1.1,) | transketolase; glycolaldehydetransferase [EC:2.2.1.1] | |
| PIAMOCKP_01006 | K01807 | rpiA | ribose 5-phosphate isomerase A [EC:5.3.1.6] | |
| PIAMOCKP_01091 | K01783 | rpe | ribulose-phosphate 3-epimerase [EC:5.1.3.1] | |
| PIAMOCKP_01192 | K01621 | xfp/xpk | xylulose-5-phosphate/fructose-6-phosphate phosphoketolase [EC:4.1.2.9] | |
| PIAMOCKP_01581 | K01810 | GPI | glucose-6-phosphate isomerase [EC:5.3.1.9] | EDD pathway |
| PIAMOCKP_02204 | K00036 | G6PD | glucose-6-phosphate 1-dehydrogenase [EC:1.1.1.49; 1.1.1.363] | |
| PIAMOCKP_00366 | K01057 | PGLS | 6-phosphogluconolactonase [EC:3.1.1.31] | |
| PIAMOCKP_02203 | K01690 | edd | phosphogluconate dehydratase [EC:4.2.1.12] | |
| PIAMOCKP_02202 | K01625 | eda | 2-dehydro-3-deoxyphosphogluconate aldolase / (4S)-4-hydroxy-2-oxoglutarate aldolase [EC:4.1.2.14; 4.1.3.42] | |
| PIAMOCKP_00076 | K00134 | gapA | glyceraldehyde 3-phosphate dehydrogenase (phosphorylating) [EC:1.2.1.12] | Glycolysis |
| PIAMOCKP_02658 | K00927 | pgk | phosphoglycerate kinase [EC:2.7.2.3] | |
| PIAMOCKP_02887 | K15633 | gpmI | 2,3-bisphosphoglycerate-independent phosphoglycerate mutase [EC:5.4.2.12] | |
| PIAMOCKP_00692 | K01689 | ENO | Enolase [EC:4.2.1.11] | |
| PIAMOCKP_00077 | K00873 | pyk | pyruvate kinase [EC:2.7.1.40] | |
| PIAMOCKP_00143 | K01006 | ppdK | pyruvate, orthophosphate dikinase [EC:2.7.9.1] | |
| PIAMOCKP_04372 | K01007 | pps/ppsA | pyruvate, water dikinase; phosphopyruvate synthetase [EC:2.7.9.2] | |
| PIAMOCKP_00529 | K00163 | aceE | pyruvate dehydrogenase E1 component [EC:1.2.4.1] | Pyruvate oxidation |
| PIAMOCKP_00530 | K00627 | aceF/pdhC/ DLAT | pyruvate dehydrogenase E2 component (dihydrolipoamide acetyltransferase) [EC:2.3.1.12] | |
| PIAMOCKP_01896 | K24012 | acdAB | acetate---CoA ligase (ADP-forming) [EC:6.2.1.13] | |
| PIAMOCKP_04321 | K01895 | acs | acetyl-CoA synthetase [EC:6.2.1.1] | |
| PIAMOCKP_04334 | K03737 | por/nifJ | pyruvate-ferredoxin/flavodoxin oxidoreductase [EC:1.2.7.1 1.2.7.-] | |
| PIAMOCKP_03969 | K01647 | CS/gltA | citrate synthase [EC:2.3.3.1] | TCA cycle |
| PIAMOCKP_03637 | K01682 | acnB | aconitate hydratase 2 / 2-methylisocitrate dehydratase [EC:4.2.1.3; 4.2.1.99] | |
| PIAMOCKP_02062 | K00030 | IDH3 | isocitrate dehydrogenase (NAD+) [EC:1.1.1.41] | |

| Locus tag | KEGG ID | Gene | Description | Pathway |
|---|---|---|---|---|
| PIAMOCKP_00808 | K01903 | sucC | succinyl-CoA synthetase beta subunit [EC:6.2.1.5] | |
| PIAMOCKP_00809 | K01902 | sucD | succinyl-CoA synthetase alpha subunit [EC:6.2.1.5] | |
| PIAMOCKP_02999 | K00164 | sucA/OGDH | 2-oxoglutarate dehydrogenase E1 component [EC:1.2.4.2] | |
| PIAMOCKP_03000 | K00658 | sucB/DLST | 2-oxoglutarate dehydrogenase E2 component (dihydrolipoamide succinyltransferase) [EC:2.3.1.61] | |
| PIAMOCKP_02929 | K00241 | sdhC | succinate dehydrogenase /fumarate reductase, cytochrome b subunit [EC:1.3.5.1; EC:1.3.5.4] | |
| PIAMOCKP_02930 | K00242 | sdhD | succinate dehydrogenase/fumarate reductase, membrane anchor subunit [EC:1.3.5.1; EC:1.3.5.4] | |
| PIAMOCKP_02931 | K00239 | sdhA | succinate dehydrogenase /fumarate reductase, flavoprotein subunit [EC:1.3.5.1; EC:1.3.5.4] | |
| PIAMOCKP_02932 | K00240 | sdhB | succinate dehydrogenase succinate dehydrogenase / fumarate reductase, membrane anchor subunit [EC:1.3.5.1; EC:1.3.5.4] | |
| PIAMOCKP_01862 | K01676 | fumA/fumB (E4.2.1.2A) | fumarate hydratase, class I [EC:4.2.1.2] | |
| PIAMOCKP_00058 | K00024 | mdh | malate dehydrogenase [EC:1.1.1.37] | |
| PIAMOCKP_01126 | K09011 | cimA | (R)-citramalate synthase [EC:2.3.3.21] | Propionate synthesis |
| PIAMOCKP_01123 | K01703 | leuC | 3-isopropylmalate/(R)-2-methylmalate dehydratase large subunit [EC:4.2.1.33 4.2.1.35] | |
| PIAMOCKP_01124 | K01704 | leuD | 3-isopropylmalate/(R)-2-methylmalate dehydratase small subunit [EC:4.2.1.33 4.2.1.35] | |
| PIAMOCKP_01083 | K00052 | leuB | 3-isopropylmalate dehydrogenase [EC:1.1.1.85] | |
| PIAMOCKP_01189 | K00925 | ackA | acetate kinase [EC:2.7.2.1] | |
| PIAMOCKP_02051 | K22214 | scpC | propionyl-CoA:succinyl-CoA transferase [EC:2.8.3.27] | |
| PIAMOCKP_04321 | K01895 | acs | acetyl-CoA synthetase [EC:6.2.1.1] | |
| PIAMOCKP_01896 | K24012 | acdAB | acetate-CoA ligase/acetyl-CoA synthetase (ADP-forming) [EC:6.2.1.13] | |
| PIAMOCKP_04308 | K18005 | hoxF | [NiFe] hydrogenase diaphorase moiety large subunit [EC:1.12.1.2] | $H_2$ production |
| PIAMOCKP_04309 | K18006 | hoxU | [NiFe] hydrogenase diaphorase moiety small subunit [EC:1.12.1.2] | |
| PIAMOCKP_04310 | K18007 | hoxY | NAD-reducing hydrogenase small subunit [EC:1.12.1.2] | |
| PIAMOCKP_04311 | K00436 | hoxH | NAD-reducing hydrogenase large subunit [EC:1.12.1.2] | |
| PIAMOCKP_01682 | K00362 | nirB | nitrite reductase (NADH) large subunit [EC:1.7.1.15] | Nitrogen metabolism |
| PIAMOCKP_01689 | K00363 | nirD | nitrite reductase (NADH) small subunit [EC:1.7.1.15] | |
| PIAMOCKP_01681 | K00372 | nasA | assimilatory nitrate reductase catalytic subunit [EC:1.7.99.-] | |
| PIAMOCKP_02200 | K15864 | nirS | nitrite reductase (NO-forming) / hydroxylamine reductase [EC:1.7.2.1 1.7.99.1] | |

| Locus tag | KEGG ID | Gene | Description | Pathway |
|---|---|---|---|---|
| PIAMOCKP_02161 | K04561 | norB | nitric oxide reductase subunit B [EC:1.7.2.5] | |
| PIAMOCKP_03330 | K02588 | nifH | nitrogenase iron protein [EC:1.18.6.1] | |
| PIAMOCKP_03331 | K02586 | nifD | nitrogenase molybdenum-iron protein alpha chain [EC:1.18.6.1] | |
| PIAMOCKP_03332 | K02591 | nifK | nitrite reductase, nitrogenase molybdenum-iron protein beta chain [EC:1.18.6.1] | |
| PIAMOCKP_00171 | K00380 | cysJ | sulfite reductase (NADPH) flavoprotein alpha-component [EC:1.8.1.2] | Assimilatory sulfate reduction |
| PIAMOCKP_00987 | K00860 | cysC | adenylylsulfate kinase [EC:2.7.1.25] | |
| PIAMOCKP_02426 | K00955 | cysNC | bifunctional enzyme cysN/cysC [EC:2.7.7.4; 2.7.1.25] | |
| PIAMOCKP_02426 | K00956 | cysN | sulfate adenylyltransferase subunit 1 [EC:2.7.7.4] | |
| PIAMOCKP_02915 | K00957 | cysD | sulfate adenylyltransferase subunit 2 [EC:2.7.7.4] | |
| PIAMOCKP_03010 | K00381 | cysI | sulfite reductase (NADPH) hemoprotein beta-component [EC:1.8.1.2] | |
| PIAMOCKP_04382 | K00390 | cysH | phosphoadenosine phosphosulfate reductase [EC:1.8.4.8; 1.8.4.10] | |

**Table S5.** Key genes encoding enzymes involved in methane oxidization to $CO_2$, formaldehyde assimilation via ribulose monophosphate pathway (RuMP) and fermentation in *Methylobacter* sp. S3L5C and several metagenome-assembled genomes (MAGs) affiliated to *Methylobacter* in environment. Details of enzymes are provided in Table S4.

| | $CH_4$ oxidation | Acetate and propionate production | Malate production | Succinate production | $H_2$ production |
|---|---|---|---|---|---|
| Strain S3L5C Lake Lovojärvi (Lammi Finland) | pmoCAB, mmoXYBZDC, xoxF, mxaFI, mxaDJGACKL, fae, mtdAB, mch, ftr, fwdBCA, fdoG, fdsD, hxlAB, pfp, fba, tkt, rpiA, rpe, xfp | acs, acdAB, ackA, cimA, leuBCD | mdh, E4.2.1.2.1A | sucCD, sdhCDAB | hoxFUYH |
| MAG bin-1608 Lake Lovojärvi (Lammi Finland) | pmoBAC, xoxF, mxaJD, fae, mtdBA, mch, ftr, fwdCAB, fdoG, fdsD, hxlBA, pfp, fba, tkt, rpiA, rpe | acs, acdAB, cimA, leuDCB | mdh, E4.2.1.2.1A | sucDC, sdhBADC, | hoxHFUY |
| MAG bin-1620 Lake Lovojärvi (Lammi Finland) | pmoBAC, xoxF, mxaJD, fae, mtdAB, mch, ftr, fwdCAB, fdoG, fdsD, cynT, hxlAB, pfp, fba, tkt, rpiA, rpe, xfp | acs, acdAB, ackA, cimA, leuDCB | mdh, E4.2.1.2.1A | sucCD, sdhDCAB | hoxUF |
| MAG bin-1610 Lake Valkea Kotinen (Evo Finland) | pmoBAC, xoxF, mxaJD, fae, mtdBA, mch, ftr, fwdCAB, fdoG, fdsD, hxlBA, pfp, fba, tkt, rpiA, rpe, xfp | acs, acdAB, ackA, cimA, leuCDB | mdh, E4.2.1.2.1A | sucDC, sdhBADC | hoxHYUF |
| MAG bin-1070 Lake Keskinen Rajajarvi (Evo Finland) | pmoCB, xoxF, mxaDJ, fae, mtdAB, mch, ftr, fwdCAB, fdoG, fdsD, hxlAB, pfp, fba, tkt, rpiA, rpe | acs, acdAB, cimA, leuBCD | mdh, E4.2.1.2.1A | sucDC, sdhCDAB | hoxFUYH |
| MAG bin-113 Lake Mekkojarvi (Evo Finland) | pmoBAC, xoxF, mxaDJ, fae, mtdBA, mch, ftr, fwdBAC, fdoG, fdsD, hxlBA, pfp, fba, tkt, rpiA, rpe | acs, acdAB, cimA, leuBCD | mdh, E4.2.1.2.1A | sucDC, sdhBADC | hoxHFUY |
| MAG bin-8093 Lake Alinen Mustajarvi (Evo Finland) | pmoCAB, xoxF, mxaJD, fae, mtdBA, mch, ftr, fwdCBA, fdoG, fdsD, hxlBA, pfp, fba, tkt, rpiA, rpe, xfp | acs, acdAB, ackA, cimA, leuBDC | mdh, E4.2.1.2.1A | sucCD, sdhBADC | hoxHYUF |
| MAG bin-692 Lake Lomtjarnan (Jamtland Sweden) | pmoCB, xoxF, mxaD, fae, mtdBA, mch, ftr, fwdBCA, fdsD, hxlAB, pfp, fba, tkt, rpiA, rpe, xfp | acs, acdAB, ackA, cimA, leuCDB | E4.2.1.2.1A | sucDC, sdhCD | hoxYUFH |
| MAG bin-0689 Lake Bengtsgolen (Linkoping Sweden) | pmoCAB, mmoCDZBYX, mxaIGJFD, fae, mtdAB, mch, ftr, fwdBC, fdoG, fdsD, hxlBA, pfp, fba, tkt, rpiA, rpe, xfp | acdAB, cimA, leuCD | mdh, E4.2.1.2.1A | sdhB | hoxFUYH |
| MAG bin-0461 Lake Bjorntjarnan (Umea Sweden) | pmoCAB, xoxF, mxaDJ, fae, mtdBA, mch, fwdB, fdsD, hxlBA, pfp, fba, tkt, rpiA, xfp | acs, ackA, cimA, leuCDB | mdh, E4.2.1.2.1A | sucCD, sdhCDAB | hoxYUFH |

| | | | | | |
|---|---|---|---|---|---|
| MAG bin-0517 Lake Erken (Uppsala Sweden) | pmoBAC, xoxF, mxaDJ, fae, mtdBA, mch, ftr, fwdBCA, fdoG, fdsD, hxlAB, pfp, fba, tkt, rpiA, rpe, | acdAB, cimA, leuBCD | mdh, E4.2.1.2.1A | sucDC, sdhCDAB | hoxHYUF |
| MAG bin-840 Lake Loclat (Neuchatel Switzerland) | pmoBAC, xoxF, mxaDJ, fae, mtdBA, mch, ftr, fwdBAC, fdoG, hxlBA, pfp, fba, tkt, rpiA, rpe | acs, acdAB, cimA, leuCDB | mdh, E4.2.1.2.1A | sucDC, sdhBA | hoxFUH |
| MAG NSO1-1 Old Woman Creek Wetland (USA) | pmoCAB, mxaD, fwdB, fdsD, hxlAB | cimA | mdh | sucDC | |
| MAG *Methylobacter* sp. KS41 acidic soil bioreactor | pmoCAB (pxmABC), xoxF, mxaDJ, fae, mtdBA, mch, ftr, fwdBAC, fdoG, fdsD, hxlBA, pfp, fba, tkt, rpiA, rpe, xfp | acs, acdAB, ackA, cimA, leuBCD | mdh, E4.2.1.2.1A | sucCD, sdhBADC | hoxFUYH |
| MAG bin-63 Lacamas Lake (Washington USA) | pmoCAB, xoxF, mxaJD, fae, mtdAB, mch, ftr, fwdCAB, fdoG, fdsD, hxlAB, pfp, fba, tkt, rpiA, rpe | acs, acdAB, cimA, leuBCD | mdh, E4.2.1.2.1A | sucCD, sdhCDAB | hoxHYUF |
| MAG bin-1219 pond C5 (Kuujjuarapik-Whapmagoostui Canada) | pmoCAB, mxaD, fae, mtdA, mch, ftr, fwdCAB, fdoG, fdsD, hxlBA, pfp, fba, rpe, xfp | acs, acdAB, ackA, cimA, leuBCD | mdh, E4.2.1.2.1A | sucDC, sdhBAD | hoxHYUF |
| MAG bin-470 pond SAS2C (Kuujjuarapik-Whapmagoostui Canada) | pmoB, mxaD, fae, mtdAB, mch, fwdAB, fdoG, fdsD, hxlBA, pfp, fba, tkt, rpiA, rpe | acs, acdAB, cimA, leuBC | mdh, E4.2.1.2.1A | sucDC, sdhBADC | hoxHYU |

**Table S6.** The consumption amounts and utilization efficiencies (UE) of $CH_4$ and $O_2$ on day 33 of the strain S3L5C incubation with and the control without 20% $CH_4$ + 80% air replenishment.

| | **with $CH_4$+air replenishment** | **without $CH_4$+air replenishment** |
|---|---|---|
| Total fed $O_2$ (mM) | 16.25 ± 0.23 | 8.12 ± 0.07 |
| Total fed $CH_4$ (mM) | 17.57 ± 0.71 | 11.44 ± 0.37 |
| Consumed $O_2$ (mM) | 9.83 ± 0.81 | 5.84 ± 0.05 |
| Consumed $CH_4$ (mM) | 12.16 ± 0.72 | 7.33 ± 0.15 |
| Produced $CO_2$ (mM) | 5.18 ± 0.60 | 2.65 ± 0.02 |
| % $O_2$ UE | 60.5 ± 5.6 | 72.0 ± 1.2 |
| % $CH_4$ UE | 69.3 ± 5.0 | 64.1 ± 2.0 |

**Table S7.** Carbon mass balance and distribution of the produced organic acids-carbon mass for consumed C-$CH_4$ on day 33 of the incubation of strain S3L5C with and the control without $CH_4$+air replenishment. The headspace was replenished with 20% $CH_4$ and 80% air on day 20.

| | **with $CH_4$+air replenishment** | **without $CH_4$+air replenishment** |
|---|---|---|
| Consumed C-$CH_4$ (µmol) | 1280 ± 80.6 | 725.7 ± 16.1 |
| Produced C-$CO_2$ (µmol) | 594.0 ± 70.9 | 306.5 ± 2.78 |
| **Produced organic acid (µmol)** | | |
| Acetate | 15.62 ± 2.92 | 4.46 ± 1.80 |
| Formate | 0.1096 ± 0.18 | 0.6026 ± 0.44 |
| Malate | 0.0915 ± 0.008 | 0.0569 ± 0.007 |
| Propionate | 0.2567 ±0.053 | ND |
| **Produced organic acid (µmol C)** | | |
| Acetate | 31.24 ± 5.85 | 8.929 ±3.60 |
| Formate | 0.1096 ± 0.18 | 0.6026 ± 0.44 |
| Malate | 0.3661 ± 0.030 | 0.2277 ± 0.030 |
| Propionate | 0.7701 ± 0.16 | ND |
| Total OA | 32.48 ± 5.94 | 9.759 ±3.42 |
| **Carbon yields of byproducts converted from the consumed C-$CH_4$ (%)** | | |
| Acetate | 2.45 ± 0.49 | 1.22 ± 0.48 |
| Formate | 0.009 ± 0.015 | 0.084 ± 0.063 |
| Malate | 0.029 ± 0.004 | 0.031 ± 0.004 |
| Propionate | 0.060 ± 0.009 | ND |
| Total C-organic acids | 2.54 ± 0.50 | 1.34 ± 0.46 |
| C-$CO_2$ | 46.35 ± 3.60 | 42.24 ± 0.61 |
| Estimated C-biomass | 51.10 ± 3.99 | 56.41 ± 0.77 |

**Note:** ND, not detected; standard deviations were calculated from triplicate samples

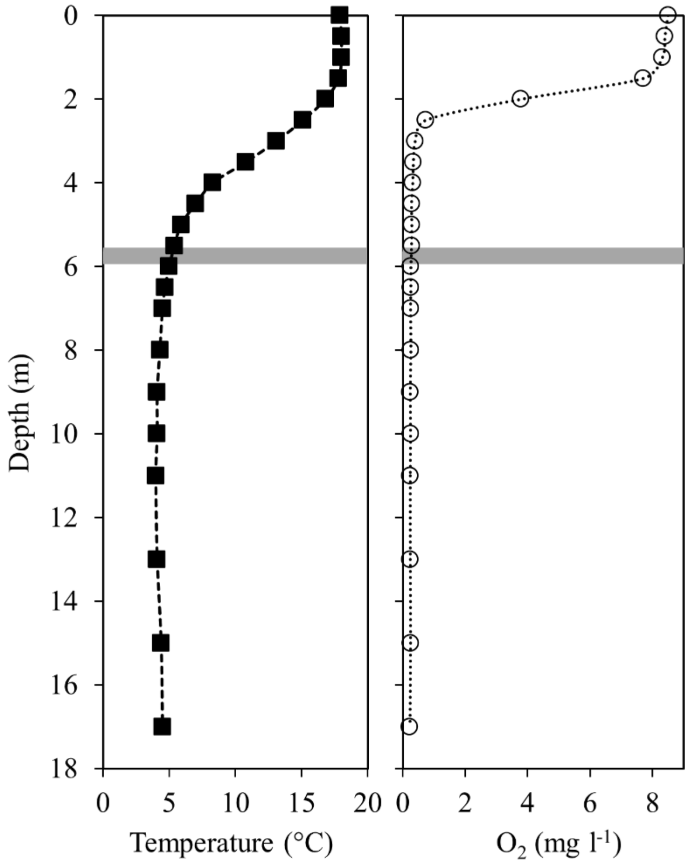


| | Sampling point 1 | Sampling point 2 |
|---|---|---|
| Depth (m) | 3.25 | 5.75 |
| N/$NH_4^+$ (µg $l^{-1}$) | 10 | 110 |
| P/$PO_4^{2-}$ (µg $l^{-1}$) | 4 | 32 |
| N/$NO_2^-$+$NO_3^-$ (µg $l^{-1}$) | 14 | 8 |
| pH | 6.83 | 6.5 |
| Temperature (°C) | 10.8-13.1 | 5.0-5.4 |
| $O_2$ (mg $l^{-1}$) | 0.34-0.40 | 0.26-0.28† |

**Fig. S1.** Water column depth profiles of temperature and dissolved oxygen (DO) of Lake Lovojärvi and physicochemical characteristics of sampling points (on September 3, 2019). Gray highlights represent lake sample used for strain S3L5C isolation. † The $O_2$-measuring instrument (ProODO) has its lowest detection limit at around 0.3 mg $l^{-1}$. Thus, the actual $O_2$ concentration is not known.

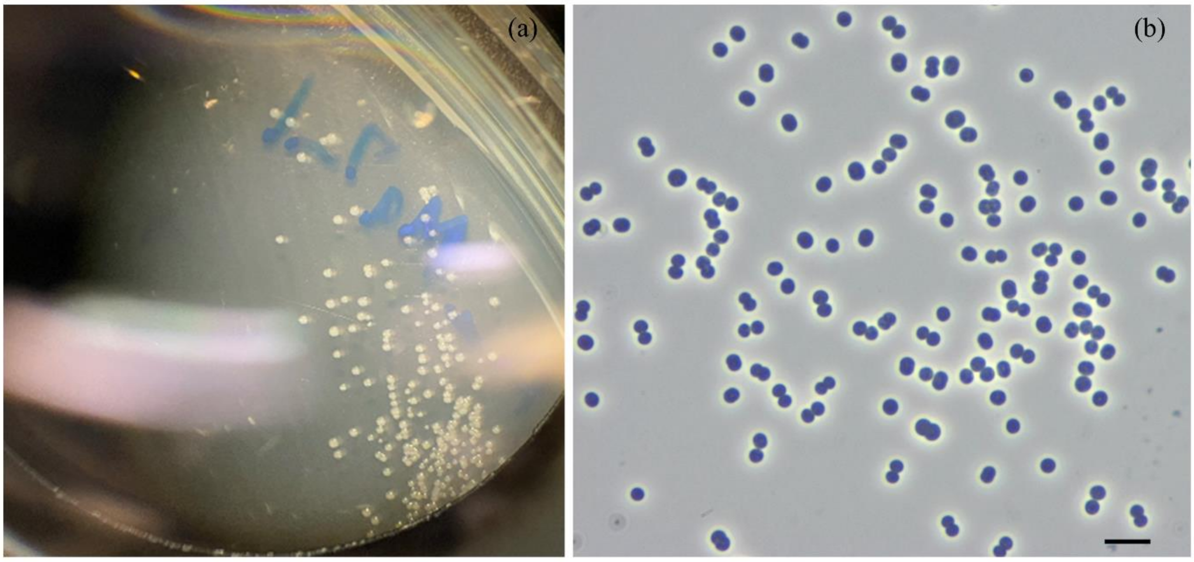


**Fig. S2.** (a) Photograph of S3L5C colonies with the size of < 0.1 mm diameter under a stereo microscope. (b) Phase-contrast light micrograph of cells of strain S3L5C; bar, 5 μm.

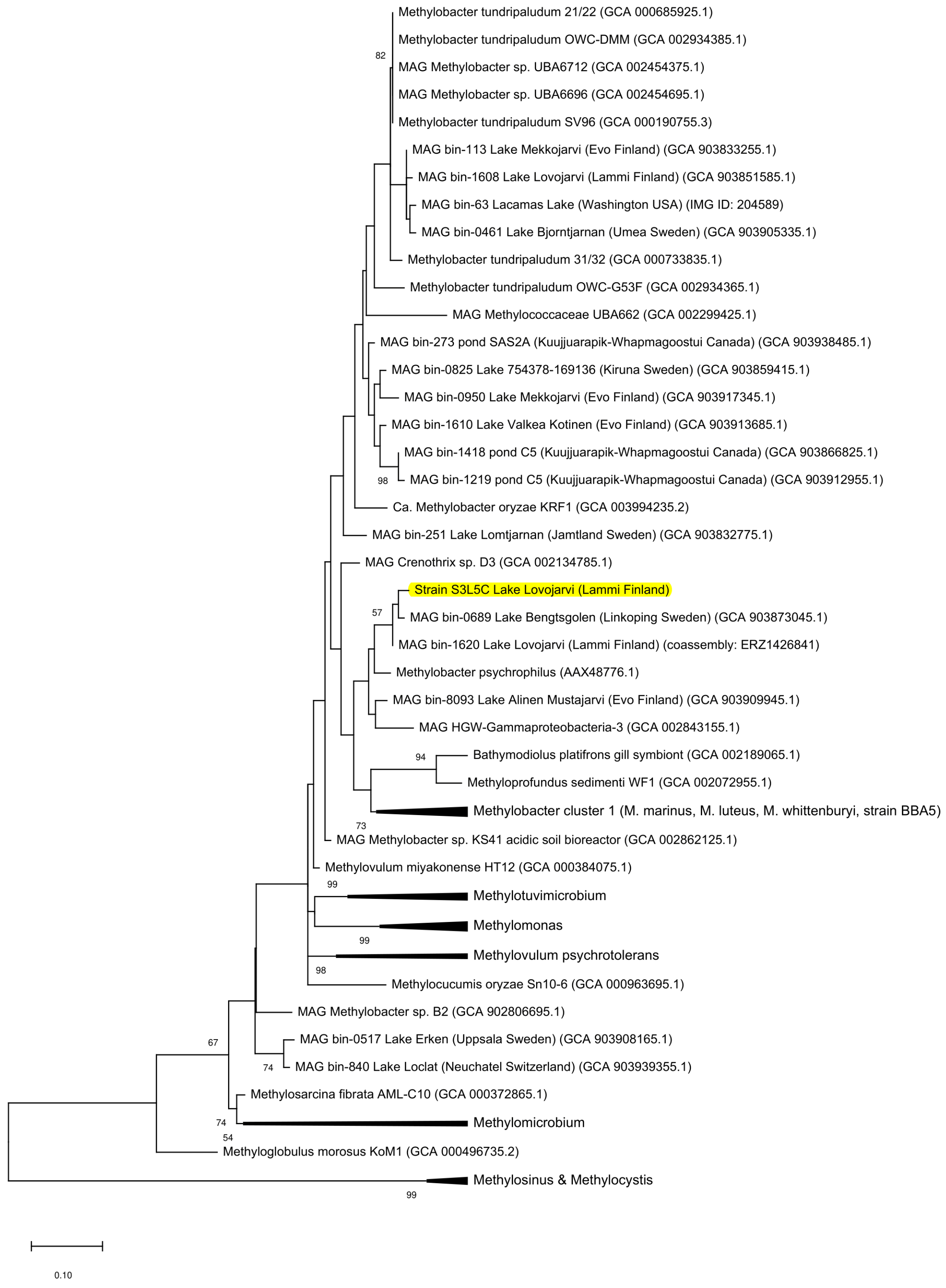


**Fig. S3.** Phylogenetic tree based on pmoA beta subunit genes of strain S3L5C (highlighted in yellow) in comparison with other pure culture methanotrophic bacteria and metagenome-assembled genomes (MAG). GenBank accession numbers are given in parentheses and bar shows 10% sequence divergence.

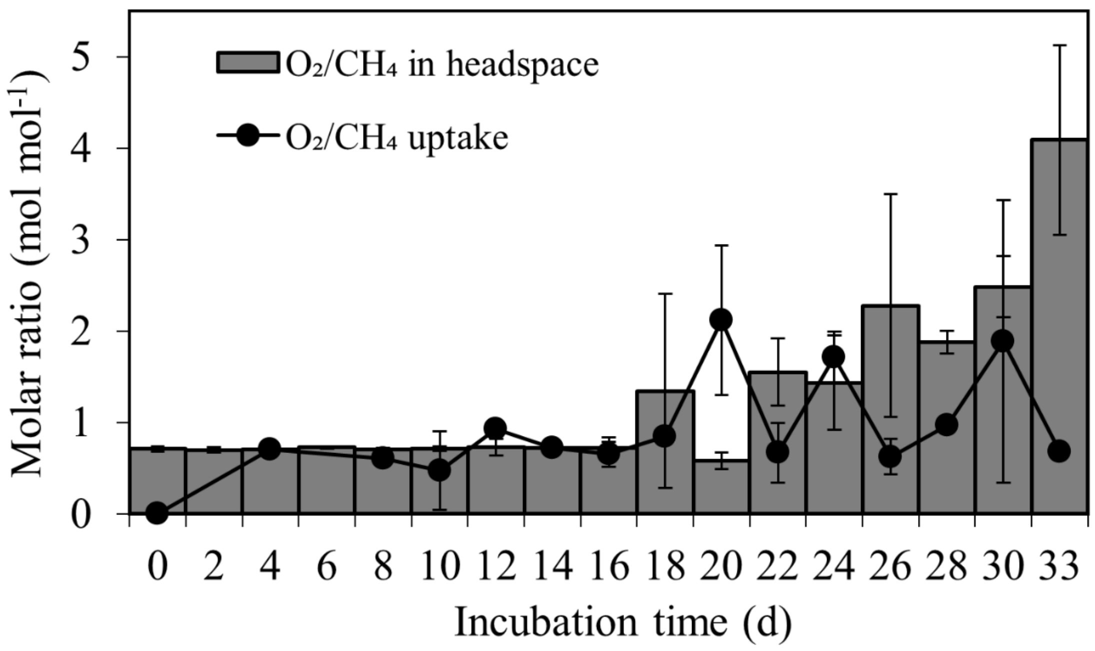


**Fig. S4.** Profiles of oxygen-to-methane ($O_2/CH_4$) molar ratio in headspace and uptake during 33-day incubation in the batch tests with 20% $CH_4$ and 80% air replenishment on day 20.

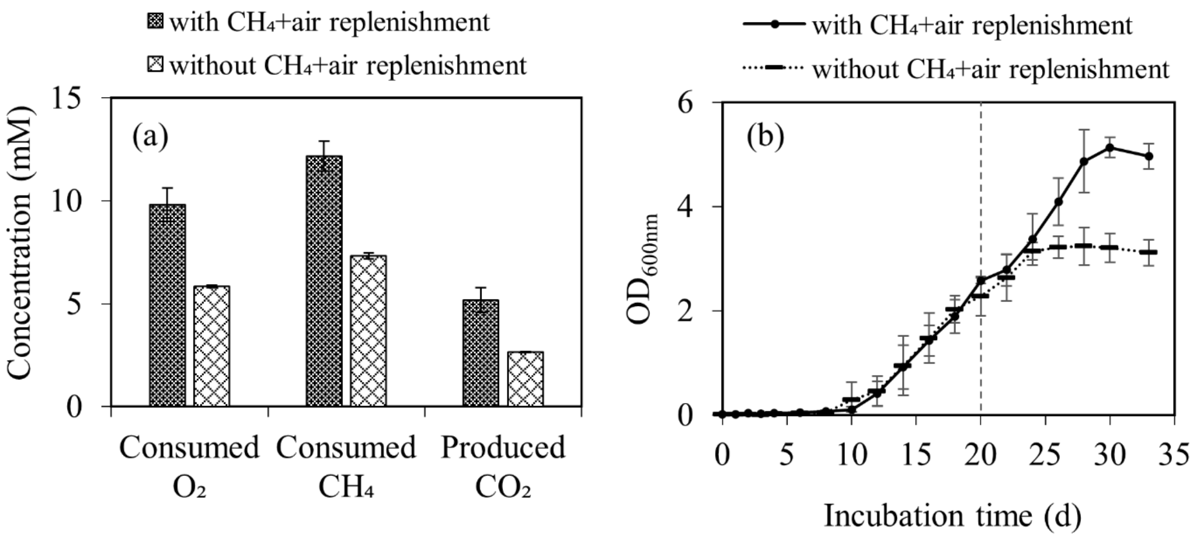


**Fig. S5.** (a) The consumed $O_2$ and $CH_4$ and produced $CO_2$ at the end of the test (day 33) and (b) biomass growth of strain S3L5C during 33-day incubation between the tests with 20% $CH_4$ and 80% air replenishment on day 20 and the control without the replenishment.
